# High-resolution transcriptomic and epigenetic profiling identifies novel regulators of COPD phenotypes in human lung fibroblasts

**DOI:** 10.1101/2022.03.28.486023

**Authors:** Uwe Schwartz, Maria Llamazares Prada, Stephanie T. Pohl, Mandy Richter, Raluca Tamas, Michael Schuler, Corinna Keller, Vedrana Mijosek, Thomas Muley, Marc A. Schneider, Karsten Quast, Joschka Hey, Claus P. Heußel, Arne Warth, Hauke Winter, Özdemirhan Serçin, Harry Karmouty-Quintana, Felix Herth, Ina Koch, Giuseppe Petrosino, Balca R. Mardin, Dieter Weichenhan, Tomasz P. Jurkowski, Charles D. Imbusch, Benedikt Brors, Vladimir Benes, Brigit Jung, David Wyatt, Heiko Stahl, Christoph Plass, Renata Z. Jurkowska

**Affiliations:** BioMed X Institute, Heidelberg, Germany; NGS Analysis Center Biology and Pre-Clinical Medicine, University of Regensburg, Regensburg, Germany; Division of Cancer Epigenomics, German Cancer Research Center (DKFZ) and Translational Lung Research Center, Member of the German Center for Lung Research (DZL), Heidelberg, Germany, Heidelberg, Germany; Division of Biomedicine, School of Biosciences, Cardiff University, Cardiff, UK; Drug Discovery Sciences, Boehringer Ingelheim Pharma GmbH & Co. KG, Germany; Immunology and Respiratory Disease Research, Boehringer Ingelheim Pharma GmbH & Co. KG – Biberach, Germany; Translational Research Unit, Thoraxklinik, University Hospital Heidelberg, University of Regensburg Germany; Translational Lung Research Center (TLRC), Member of the German Center for Lung Research (DZL), Heidelberg, Germany; Global Computational Biology and Digital Sciences, Boehringer Ingelheim Pharma GmbH & Co. KG, Germany; Ruprecht Karl University of Heidelberg, Heidelberg, Germany; Diagnostic and Interventional Radiology with Nuclear Medicine, Thoraxklinik, University of Heidelberg, Heidelberg, Germany; Diagnostic and Interventional Radiology, University Hospital Heidelberg, Heidelberg, Germany; Pathological Institute, University Hospital Heidelberg, Heidelberg, Germany; Dept. of Surgery, Thoraxklinik, University Hospital Heidelberg, Heidelberg, Germany; Department of Biochemistry and Molecular Biology, McGovern Medical School, University of Texas Health Science Center at Houston, Houston, USA; Department of Pneumology and Critical Care Medicine and Translational Research Unit, Thoraxklinik, University Hospital Heidelberg, Heidelberg, Germany; Asklepios Biobank for Lung Diseases, Department of Thoracic Surgery, Asklepios Fachkliniken München-Gauting, German Center for Lung Research (DZL), Munich, Germany; Division of Molecular Biology, School of Biosciences, Cardiff, UK; Division of Applied Bioinformatics, German Cancer Research Center, Germany; Genome Biology Unit, European Molecular Biology Laboratory (EMBL), Heidelberg, Germany; Biotherapeutics Discovery, Boehringer Ingelheim Pharma GmbH & Co. KG, Germany

**Keywords:** COPD, human lung fibroblasts, epigenome, DNA methylation, WGBS, RNA sequencing, phenotypic assays, biomarkers

## Abstract

Patients with chronic obstructive pulmonary disease (COPD) are still waiting for curative treatments. Considering the environmental cause of COPD (e.g., cigarette smoke) and disease phenotypes, including stem-cell senescence and impaired differentiation, we hypothesized that COPD will be associated with altered epigenetic signaling in lung cells. We generated genome-wide DNA methylation maps at single CpG resolution of primary human lung fibroblasts (HLFs) isolated from distal parenchyma of ex-smoker controls and COPD patients, with both mild and severe disease. The epigenetic landscape is markedly changed in lung fibroblasts across COPD stages, with DNA methylation changes occurring predominantly in regulatory regions, including promoters and enhancers. RNA sequencing of matched fibroblasts demonstrated dysregulation of genes involved in proliferation, DNA repair, and extracellular matrix organization. Notably, we identified epigenetic and transcriptional dysregulation already in mild COPD patients, providing unique insights into early disease. Integration of profiling data identified 110 candidate regulators of disease phenotypes, including epigenetic factors. Using phenotypic screens, we verified the regulator capacity of multiple candidates and linked them to repair processes in the human lung.

Our study provides first integrative high-resolution epigenetic and transcriptomic maps of human lung fibroblasts across stages of COPD. We reveal novel transcriptomic and epigenetic signatures associated with COPD onset and progression and identify new candidate regulators involved in the pathogenesis of chronic respiratory diseases. The presence of various epigenetic factors among the candidates demonstrates that epigenetic regulation in COPD is an exciting research field that holds promise for novel therapeutic avenues for patients.

## Introduction

Chronic obstructive pulmonary disease (COPD) is a prevalent, smoke-related disease characterized by persistent inflammation of the lung epithelium, irreversible airway remodeling, and destruction of the alveolar tissue (emphysema) (Barnes et al. 2015; Rabe and Watz 2017; GOLD 2021). COPD- related mortality is increasing and it already affects more than 3 million people worldwide every year (WHO 2019). However, despite its prevalence, there is currently no treatment to halt progression of COPD, as none of the existing drugs can modify the long-term decline in lung function. COPD is a heterogeneous disease with variable clinical manifestations and responses to therapy where patient stratification remains challenging (Woodruff et al. 2015; Agusti et al. 2017; Garudadri and Woodruff 2018; Barnes 2019a).

Fibroblasts are ubiquitous mesenchymal cells found in the parenchyma and the outer layer of airways and vessels in the adult lung (Phan 2008). They have essential functions in lung homeostasis, maintenance of stem cells, wound healing, and tissue repair. In COPD, airway fibroblasts are the key cells contributing to the excessive deposition of extracellular matrix, small-airway fibrosis and airway remodelling (Barnes 2019b). In turn, parenchymal fibroblasts from patients with COPD/emphysema show reduced proliferation (Nobukuni et al. 2002; Holz et al. 2004), contractility and migration *in vitro* (Togo et al. 2008), are senescent (Muller et al. 2006), display altered growth factor response (Noordhoek et al. 2003; Togo et al. 2008) and express increased levels of pro-inflammatory cytokines (Zhang et al. 2012), indicative of a reduced tissue-repair capacity. The altered function of alveolar fibroblasts also contributes to epithelial progenitor dysfunction, establishing fibroblasts as a critical cell type contributing to the development of emphysema (Plantier et al. 2007; Kulkarni et al. 2016). However, it remains unknown how these phenotypic changes in parenchymal fibroblasts are encoded at the molecular level.

Numerous genetic loci have been associated with COPD and lung function (Wilk et al. 2009; Hancock et al. 2010; Soler Artigas et al. 2011; Cho et al. 2014; Wain et al. 2015; Hobbs et al. 2017; Wyss et al. 2018; Sakornsakolpat et al. 2019), yet they explain only a small fraction of COPD risk. Transcriptional programs in cells are regulated by a landscape of epigenetic modifications that modulate chromatin structure and thereby control gene expression. Smoking is the most prominent risk factor for COPD, and its impact on epigenetic landscape remodeling is well established (Belinsky et al. 2002; Chen et al. 2013; Zeilinger et al. 2013; Wan et al. 2015). Earlier studies also provided strong evidence for the association of dysregulated DNA methylation and COPD in blood (Qiu et al. 2012; Busch et al. 2016; Carmona et al. 2018), sputum (Sood et al. 2010), oral mucosa (Wan et al. 2015), lung tissue (Sood et al. 2010; Yoo et al. 2015; Morrow et al. 2016; Sundar et al. 2017), bronchial brushings (Vucic et al. 2014), fibroblasts (Clifford et al. 2018) and macrophages from a mouse model of muco-obstructive disease (Hey et al. 2021). Notably, DNA methylation changes were associated with altered expression of genes and pathways important to COPD pathology. However, these studies were either performed on material encompassing mixed cell populations or/and used low-resolution approaches and could therefore not resolve differential gene expression and methylation changes caused by a specific cell type during COPD development and progression. To date, the full epigenomic landscape of purified COPD cells remains uncharted and thus, the precise epigenetic changes and their contribution to altered transcriptional patterns in COPD are still unknown.

To identify the epigenetic and functional alterations associated with COPD in parenchymal fibroblasts, we used whole-genome bisulfite sequencing to profile DNA methylation and RNA sequencing to measure gene expression changes in primary fibroblasts from patients with COPD and matched ex-smoker controls. Importantly, we hypothesized that epigenetic modifications would arise early during COPD development, thus, we analyzed cells from patients at different COPD stages. Our data provide integrative epigenetic and transcriptomic maps of fibroblasts at high resolution. It reveals pathways and novel candidate regulators, including epigenetic factors, that might be involved in the pathogenesis of chronic respiratory diseases.

## Results

### Genome-wide epigenetic changes occur early in primary lung fibroblasts during COPD

To assess the extent of epigenetic remodeling in COPD genome-wide, we generated high-resolution DNA methylomes of primary lung fibroblasts isolated from well-matched control donors (no COPD, n=3) and patients with established COPD (stage II-IV according to Global Initiative for Chronic Obstructive Lung Disease (GOLD) (GOLD 2021), n=5), which can be classified based on lung function (**Figure 1A**, **Suppl. Fig. 1A-B**). Isolated cells displayed a typical fibroblast morphology, and their high purity was confirmed by fluorescence-activated cell sorting (FACS) and immunofluorescence (IF) staining (**Suppl. Fig. 1C-D**). We used tagmentation-based whole-genome bisulfite sequencing (T-WGBS) for DNA methylation profiling, allowing genome-scale assessment of DNA methylation at single CpG resolution from low cell numbers (Wang et al. 2013) (**Figure 1B**).

**Figure 1.**
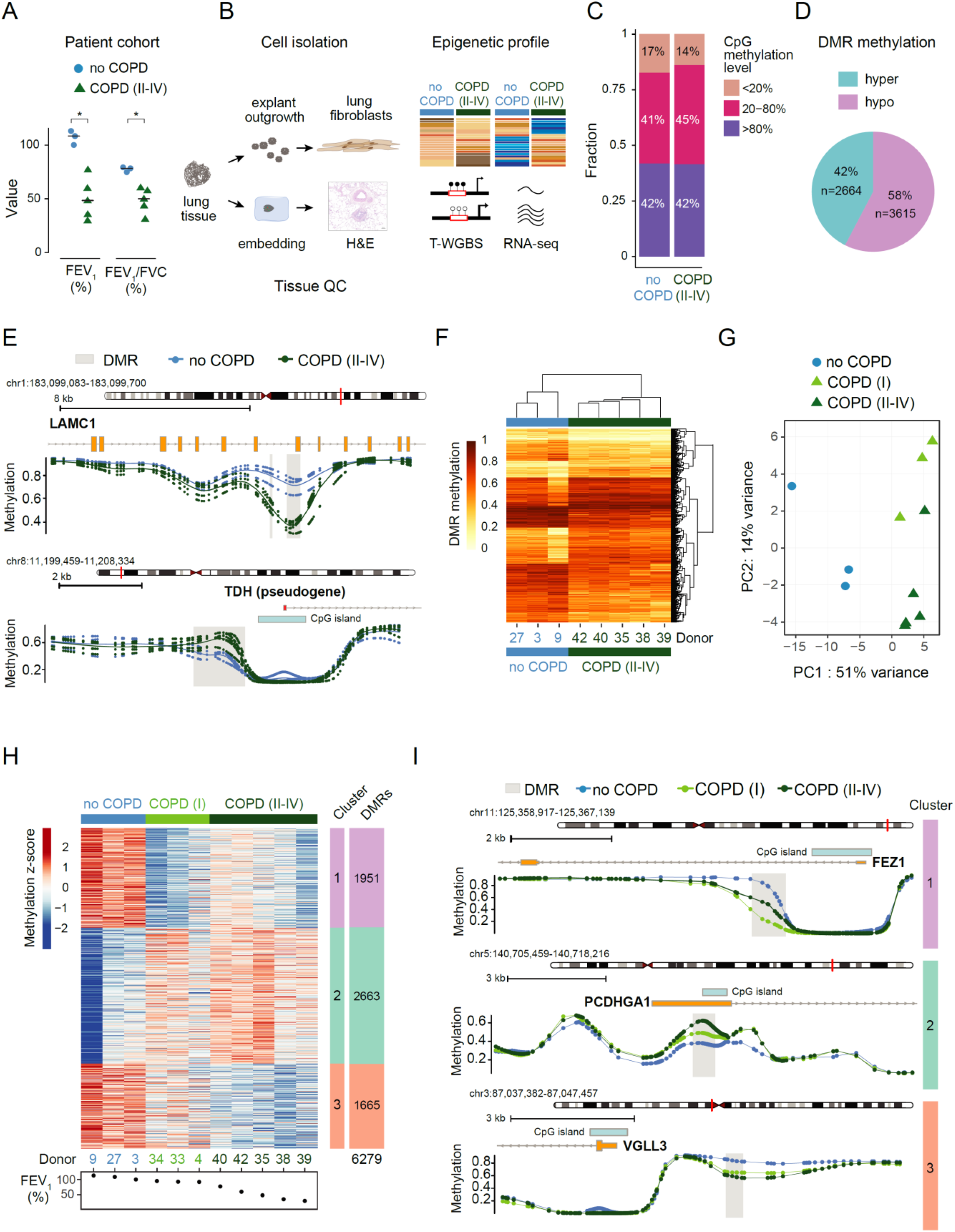
Genome-wide DNA methylation changes occur early in human lung fibroblasts during COPD and progress with disease development. **A** Lung function data of COPD (II-IV) and no COPD (ex-smoker control group) donors used in this study. Lung function between the two groups is significantly different (*p-value<0.05, unpaired non-parametric Mann-Whitney t-test). FEV_1_, forced expiratory volume in 1 s; FVC, forced vital capacity. **B** Schematic diagram illustrating the experimental approach used for epigenetic (T-WGBS) and transcriptomic (RNA-seq) profiling of purified primary parenchymal lung fibroblasts. **C-I** Data of tagmentation-based WGBS (T-WGBS) of primary fibroblasts from no COPD and COPD (II-IV) patients were analyzed on single CpGs level (**C**) and on differentially methylated regions (DMRs) (**D-I**). **C** Genome wide CpG methylation statistic. Bar plot showing the fraction of high (>80%), moderate (20-80%) and low (<20%) methylated CpGs in no COPD and COPD (II-IV) samples. **D** Number of hyper- or hypomethylated DMRs in COPD (II-IV). **E** Detailed view of a representative hypo- (top) and hypermethylated (bottom) DMR (grey box). CpG methylation levels of each individual donor (dots) and the group average (lines) methylation profile of three no COPD (blue) and five COPD (II-IV) (dark green) donors are displayed. RefSeq annotated genes and CpG islands are indicated. **F** Heatmap of 6,279 DMRs identified in COPD (II-IV). Statistically significant DMRs (at significance level=0.1; see methods for DMR calling details) with at least three CpGs and a mean difference in methylation between no COPD and COPD (II-IV) of ≥ 10% were selected. Color shades indicate low (light) or high (dark) DMR methylation. **G** Principal component analysis (PCA) of COPD (II-IV) (dark green), no COPD (blue) and mild COPD (I) (light green, samples not used for initial DMR calling) on identified 6,279 DMRs. **H** K-means clustering of all DMRs identified between no COPD and COPD (II-IV) across all samples, including COPD (I). Three clusters were identified. Cluster 1 shows early hypomethylation in COPD (I), clusters 2 and 3, gradual hyper- and hypomethylation, respectively. Donors are sorted according to their FEV_1_ value as indicated at the bottom. **I** Representative methylation profiles at selected DMRs from each cluster. Group median CpG methylation is shown for no COPD (blue), COPD (I) (light green) and COPD (II-IV) (dark green). RefSeq annotated genes and CpG islands are indicated. FEV_1_, forced expiratory volume in 1 s.

We observed no significant differences in the global levels of methylated cytosines between COPD and control samples (**Figure 1C**), suggesting that, in contrast to cancer cells (Esteller 2008), COPD is not associated with a global drop or gain of methylation in lung fibroblasts. When we looked at local aberrant DNA methylation between no COPD and COPD (II-IV), we found numerous distinct regions. Using CpG sites covered at least 4x in all samples and a ≥10% methylation-difference cutoff, we identified 6,279 differentially methylated regions (DMRs) (p-value < 0.1, see Methods part for details) (**Figure 1D-F, Table 1**), indicating widespread methylation alterations in primary human lung fibroblasts of COPD patients. The distribution of methylation differences across DMRs demonstrated a more prominent loss of methylation, suggestive of a more permissive chromatin state in COPD (58% of DMRs, 3,615 hypomethylated regions, **Figure 1D).** The remaining 2,664 regions showed increased methylation (42% of the DMRs, 2,664 hypermethylated regions, **Figure 1D**). Called DMRs contained 8 CpG sites on average and showed a median size of 479 bp (**Suppl. Fig. 1E-F**), indicating that specific regions are altered.

To investigate whether DNA methylation changes occur early in COPD development and identify alterations associated with disease progression, we additionally performed T-WGBS on fibroblasts isolated from mild COPD patients (GOLD I, n=3), with noticeable obstruction (FEV_1_/FVC<70%) but preserved FEV_1_ (>80%, **Suppl. Fig. 1A**), and integrated them into the analysis. Notably, as demonstrated by the principal component analysis (PCA) on all 6,279 previously identified DMRs, COPD (I) samples grouped with the COPD (II-IV) samples on the first principal component (**Figure 1G**), confirming the COPD specific DMR calling using independent test samples (COPD (I) samples were not used for the initial DMR selection). Furthermore, the COPD (I) samples were separated from COPD (II-IV) on the second principal component, indicating that DNA methylation data might provide information about disease progression (**Figure 1G**). Consistent with the PCA, hierarchical clustering using the identified DMRs showed that COPD (I) samples group with COPD (II-IV) samples, demonstrating that COPD-associated methylation changes occur early in the disease pathogenesis (**Suppl. Fig. 1G**). This important discovery suggests that DNA methylation might provide a sensitive biomarker to separate early COPD patients from smokers with preserved lung function.

Since COPD is a progressive lung disease, we wanted to gain more insights into the kinetics of DNA methylation changes between the three donor groups with different disease severity (no COPD, COPD (I), COPD (II-IV)). For this, we performed k-means clustering on the 6,279 identified DMRs using all samples. This analysis defined 3 main clusters displaying DNA methylation changes that progressed with increasing disease severity denoted by the decline of lung function of patients (decreasing FEV_1_, **Figure 1H**). Cluster 1 and 3 showed loss of methylation at 1,951 DMRs and 1,665 DMRs, respectively, while cluster 2 (2,663 DMRs) displayed a progressive gain of methylation in COPD. Cluster 1 reveals regions with pronounced demethylation occurring already in COPD (I), while cluster 3 shows gradual loss of methylation as disease severity increases (**Figure 1H-I**).

To shed light on the cellular processes and pathways affected by aberrant DNA methylation changes in COPD, we linked DMRs to the nearest gene and performed gene ontology (GO) enrichment analysis using Genomic Regions Enrichment of Annotations Tool (GREAT) (McLean et al. 2010). As hypermethylated DMRs did not reveal any significant enrichment in GREAT, we focused on the hypomethylated DMRs. Among the top categories, we identified cellular response to hypoxia, regulation of focal adhesion assembly, negative regulation of transforming growth factor beta (TGFβ) receptor signaling, and epithelial to mesenchymal transition (**Suppl. Fig. 1H**). These biological processes are relevant for COPD development and progression (Konigshoff et al. 2009; Barnes et al. 2015; Rabe and Watz 2017; Barnes et al. 2019), indicating that DNA methylation changes occur near genes that are critically involved in COPD pathogenesis and might therefore contribute to disease phenotypes in lung fibroblasts. Specific examples include hypomethylation of genes implicated in negative regulation of TGFβ receptor signaling (e.g., SMAD6, SMAD7, VASN, SKI, PMEPA1, **Table 1**) (Miyazono 2000), potentially explaining the reduced response to TGFβ and decreased repair capacity of lung fibroblasts observed in emphysema (Togo et al. 2008).

In summary, DNA methylation profiling of COPD samples across disease stages demonstrates that genome-wide epigenetic changes occur already early in COPD development and many of them progress with disease development.

### DNA methylation changes occur at regulatory regions in COPD lung fibroblasts

Methylome analysis identified genome-wide changes of DNA methylation in primary HLFs of COPD patients. To better understand the functional role of aberrant methylation in COPD, we investigated the distribution of DMRs across the genome. We observed a different distribution of regions displaying loss or gain of methylation, with hypomethylated DMRs predominately located in intronic sequences and hypermethylated DMRs preferentially found in intergenic regions (**Figure 2A**). Notably, both types of DMRs were overrepresented at regulatory and gene coding sequences compared to the genomic background (**Figure 2B**), with a stronger enrichment for hypomethylated DMRs. Thus, hypomethylated DMRs are located 4 times more often at promoter sequences than expected by chance (**Figure 2B**). Further intersection with known regulatory genomic features annotated by the ENCODE Chromatin States (Ernst et al. 2011) revealed a strong enrichment of hypomethylated DMRs in active promoters and enhancers, indicating their potential regulatory role (**Figure 2C**-**D, Table 1**). The significant association with active enhancer elements was confirmed by the local increase of enhancer defining chromatin marks (H3K4me1 and H3K27ac) (Heintzman et al. 2009; Rada-Iglesias et al. 2010) in the center of the hypomethylated DMRs (**Figure 2D-E**). Conversely, the hypermethylated DMRs were overrepresented at Polycomb-repressed regions, defined by the presence of H3K27me3 (**Figure 2C and 2E**).

**Figure 2.**
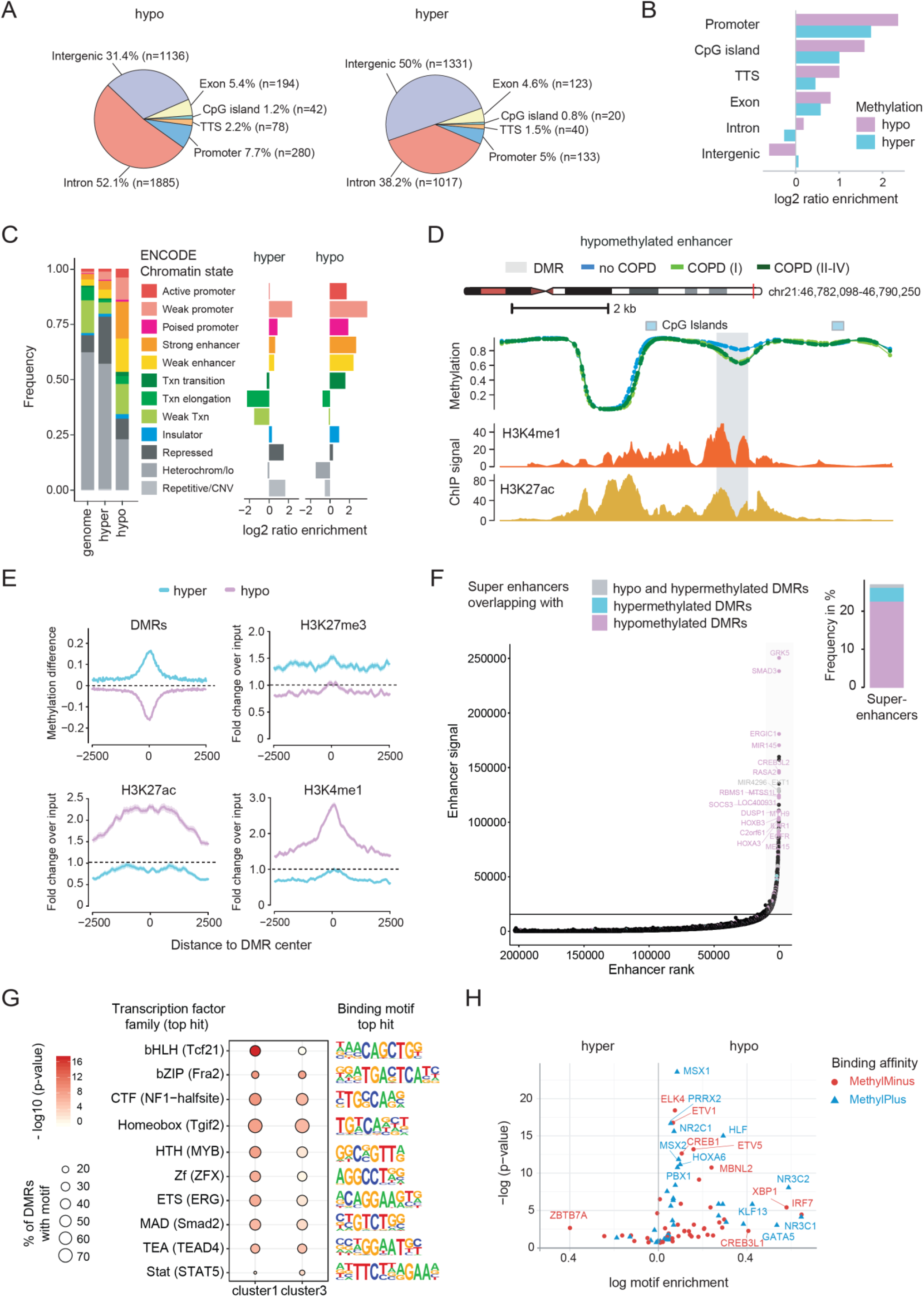
DNA methylation changes occur at regulatory regions in primary human lung fibroblasts cells during COPD. **A-B** Genomic location of identified DMRs. **A** Distribution of genomic features overlapping with hypo- (left) and hypermethylated (right) DMRs. **B** Enrichment of genomic features at hypo- (purple) and hypermethylated (cyan) DMRs compared to a sampled background of 10,000 regions exhibiting no significant change in methylation. TTS, transcription termination site. **C** Distribution of human lung fibroblast specific chromatin states (ENCODE accession: ENCFF001TDQ) at hypo- and hypermethylated DMRs. Fraction of DMRs overlapping with specific chromatin states is shown on the left panel. The genome background was sampled using 10,000 regions with matching GC content exhibiting no significant change in methylation. Chromatin state enrichment relative to the genome background is illustrated in the right panel. **D** Genome browser view of an example DMR at a putative enhancer region. Group median CpG methylation is shown for no COPD (blue), COPD (I) (light green) and COPD (II-IV) (dark green). At the bottom the level of enhancer marks is depicted as fold-change over control: H3K4me1 (ENCODE accession: ENCFF102BGI) and H3K27ac (ENCODE accession: ENCFF386FDQ). **E** Alterations of DNA methylation and selected histone marks around DMRs. Solid lines represent the mean profile and shaded lines the standard error of the mean across all summarized regions. **F** Ranking of enhancer elements, defined by the co-occurrence of H3K4me1 and H3K27ac signals in human lung fibroblasts. The horizontal line defines the signal and corresponding rank threshold used to identify super enhancers (SE). Selected SE overlapping with DMRs are annotated and the nearest gene to the SE is indicated. The fraction of SEs overlapping with hypo- (purple), hypermethylated (cyan) or both (grey) is illustrated in the bar plot on the right panel. **G** Transcription factor motifs most enriched at DMRs overlapping with strong enhancers (ENCODE chromatin states) from cluster 1 and 3 (see Figure 1H). **H** Enrichment of methylation sensitive transcription factor motifs at hypo- (right) and hypermethylated (left) DMRs. Methylation sensitive motifs were derived from the study of Yin *et al*. (Yin et al. 2017). Transcription factors, whose binding affinity was impaired upon methylation of their corresponding DNA motif are shown in red (MethylMinus) and transcription factors, whose binding affinity was increased, in blue (MethylPlus).

Since we detected enrichment of hypomethylated DMRs residing in regions broadly marked by H3K4me1 and H3K27ac (**Figure 2D-E**), we used the intensity of the H3K4me1 and H3K27ac signals in the ENCODE ChIP-seq data (Davis et al. 2018) to classify super enhancers (SE) in human lung fibroblasts. Next, we tested the overlap of the identified SE with the DMRs identified in COPD HLFs. About a quarter of all SE contained at least one DMR (**Figure 2F**, right panel, **Suppl. Fig. 2A**, examples shown in **Suppl. Fig. 2B-C**). Consistent with the chromatin state analysis, hypomethylated DMRs were preferentially associated with SE and coincided with the best scoring SE (**Figure 2F**, purple bar and labels), indicating that these SE may become differentially regulated in COPD. Finally, SE were assigned to nearby genes that they may regulate. SMAD3, GRK5, ERGIC1, CREB3L2, and RASA2 genes were associated with the most active super-enhancers overlapping with hypomethylated DMRs (**Figure 2F**, hockey plot, purple labels, and **Suppl. Fig. 2B-C**).

We conclude that methylation changes identified in COPD HLFs, especially hypomethylation, occur at regulatory regions, including strong enhancers.

### DMRs in COPD show enrichment of binding motifs for key lung transcription factors

The hypomethylated regions identified in WGBS data might reflect the binding of transcription factors and can be therefore used for foot-printing their binding sites (Stadler et al. 2011). We observed a significant enrichment of binding motifs of several transcription factors in the hypomethylated DMRs at strong enhancers, with the highest enrichment of TCF21 motif in the early DMR cluster (cluster 1, p-value: 1×10^-17^) and FOSL2/FRA2 in the progressive DMR cluster (cluster 3, p-value: 1×10^-8^) (**Figure 2G**). TCF21 mediates fibroblast fate specification in multiple organs and is required for alveolar development (Quaggin et al. 1999). In turn, FOSL2/FRA2 is a known regulator of wound repair and TGFβ-mediated fibrosis (Eferl et al. 2008). Our data establish TCF21 and FOSL2/FRA2 as potential mediators of aberrant epigenetic changes at strong enhancers in COPD fibroblasts.

Since DNA methylation can also directly interfere with the binding of transcriptional regulators to DNA (Yin et al. 2017), we performed motif analysis in the identified hypo- and hypermethylated DMRs to identify transcription factors reported to change their binding affinity upon methylation of their motifs (Yin et al. 2017). At hypomethylated DMRs, which are overrepresented at regulatory sites, numerous methylation-sensitive transcription-factor motifs were significantly enriched, suggesting either increased (methyl-minus, red dots) or attenuated (methyl-plus, blue triangles) DNA binding in COPD (**Figure 2H**). Among the transcription factors exhibiting higher binding affinities towards methylated DNA (methyl-plus, blue triangles), we identified motifs of nuclear receptors, known regulators of cellular homeostasis, development, and metabolism (**Figure 2H**). Our data suggest that their DNA binding might be abrogated at regulatory sites in COPD due to loss of methylation. A few motifs of methylation-sensitive transcription factors were enriched in the hypermethylated DMRs, consistent with their location in repressive regions of the genome. The strongest enrichment was observed for ZBTB7A, a known repressor associated with the TGFβ signaling pathway (Shen et al. 2017). ZBTB7A preferentially binds unmethylated DNA (Yin et al. 2017) (methyl-minus, **Figure 2H**) indicating that its DNA binding in COPD might be hindered due to motif hypermethylation.

In summary, our data link aberrant DNA methylation in COPD fibroblasts to imbalanced transcription factor binding and provides insights into potentially disturbed regulatory networks in COPD.

### Gene expression changes accompany epigenetic modifications in COPD

Our genome-wide DNA methylation analysis identified methylation changes at promoter and enhancer regions, suggesting that DMRs may have regulatory effects on gene expression. To assess whether epigenetic changes are associated with gene expression changes in COPD, we performed RNA-seq analysis on fibroblast samples matching those used for T-WGBS. This analysis identified 333 up-regulated and 287 down-regulated genes between no COPD (n=3) and COPD (II-IV) (n=5; FDR < 0.05 and |log2(fold-change)| > 0.5, **Figure 3A**, **Table 2**), including several long non-coding RNAs (**Figure 3A** orange labels, **Suppl. Fig. 3A**), providing a transcriptional signature of COPD. Enrichment analysis of the differentially expressed genes revealed that genes up-regulated in COPD are involved in cell cycle regulation, DNA replication and DNA repair (**Suppl. Fig. 3B**). In turn, down-regulated genes are associated with extracellular matrix (ECM) organization and cholesterol biosynthesis (**Suppl. Fig. 3C**).

**Figure 3.**
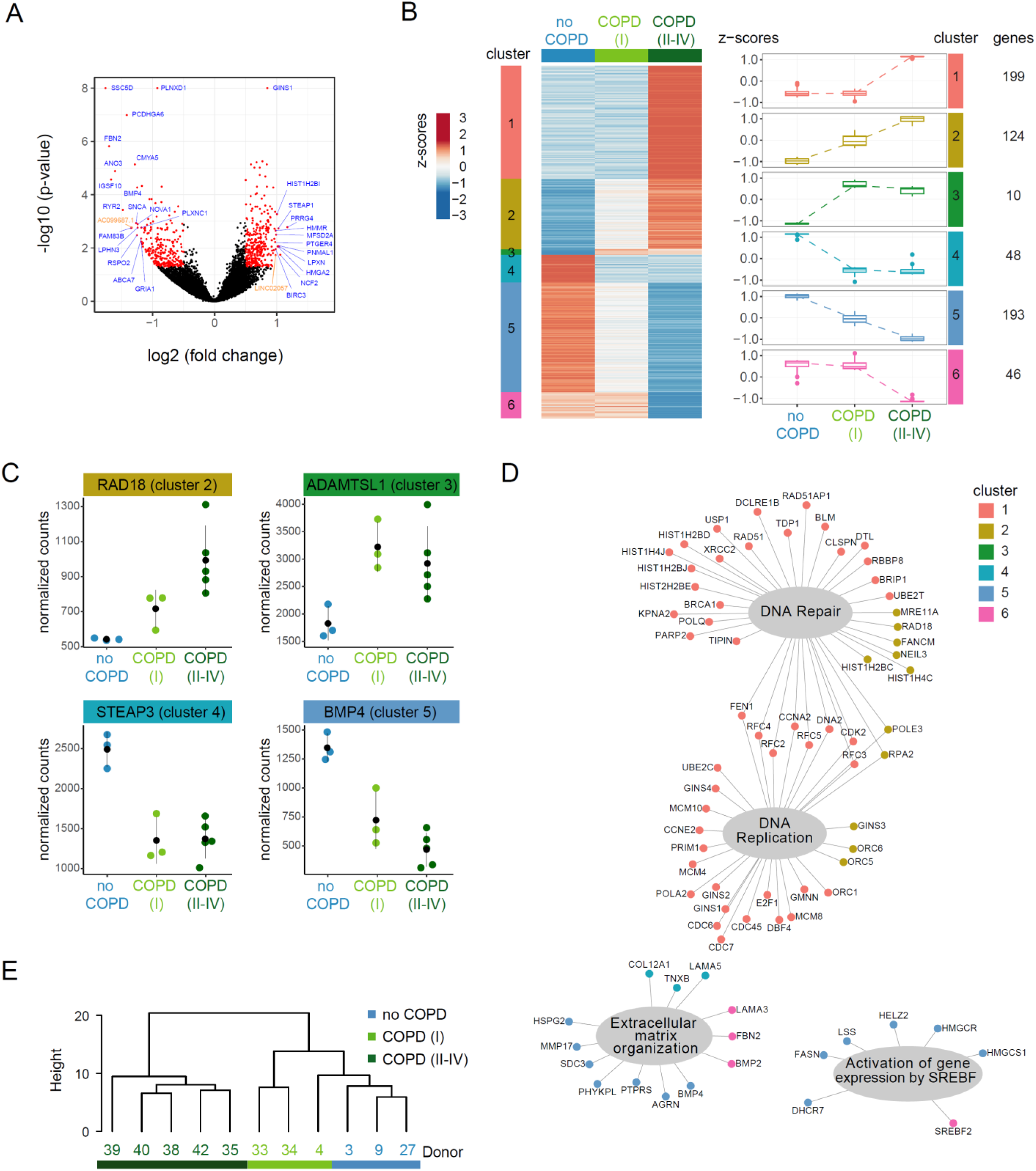
DNA methylation changes in primary human lung fibroblasts are accompanied by gene expression changes in COPD. **A** Volcano plot of differentially expressed genes (DEG) (red dots; FDR < 0.05 and |log2(fold change)| > 0.5) in COPD (II-IV) compared to no COPD controls. Protein-coding (blue) and lincRNA (orange) with the highest expression change or lowest p-values are labeled. **B** Self-organizing maps (SOM) clustering based on the scaled median expression level per group of the 620 DEG identified between COPD (II-IV) and no COPD samples. DEGs were grouped into 6 distinct clusters showing different kinetics in COPD progression: Clusters 1 and 6 show late changes. Clusters 2 and 5 display changes gradually progressing with disease severity. Clusters 3 and 4 correspond to early changes observed already in COPD (I). **C** Selected examples of DEG across disease stages from clusters 2-5. **D** DEGs associated with altered biological processes (grey bubbles) in COPD. DEG nodes are colored according to their corresponding gene expression kinetic in COPD (clusters defined in **3B**). **E** Unsupervised hierarchical clustering of all samples, including COPD (I) based on DEGs (n=620) identified between no COPD and COPD (II-IV) samples.

As COPD (I) samples (n=3) were not used to identify differentially expressed genes (DEGs), we used them as an independent test-set to validate the obtained results and gain insights into gene expression kinetics in disease progression. Hierarchical clustering and PCA of all samples on the identified DEGs or 500 most variable genes, respectively, revealed that COPD (I) samples cluster together and show higher similarity to the no COPD group (**Figure 3E**, **Suppl. Fig. 3D**). To further resolve gene expression signatures of COPD states, we performed self-organizing map (SOM) clustering (Wehrens and Kruisselbrink 2018) using all DEGs (n=620). We identified 6 clusters showing different kinetics related to COPD progression (**Figure 3B-D, Table 2**). Clusters 1 and 6 encompass genes whose expression is not yet changed in COPD (I) but gets dysregulated at later stages of COPD development (**Figure 3B**). Here, multiple genes involved in DNA replication (e.g., MCM10, ORC5, GINS3) or DNA double-strand break repair (e.g., BRCA1, FANCM, USP1, RAD51) are present (**Figure 3D**). Clusters 2, 3, 4, and 5 feature gene subsets already dysregulated in COPD (I) and may serve as early disease markers (**Figure 3B**, examples displayed in **Figure 3C)**. Notably, identification of early gene expression changes associated with COPD development is of high clinical relevance, as it might offer a unique advantage for disease-modifying therapies.

To assess the extent of epigenetic remodeling in COPD, we analyzed the expression changes of epigenetic enzymes and readers(Medvedeva et al. 2015). 38 epigenetic factors were differentially expressed, with the majority (76%) showing up-regulation in COPD (**Table 2**). Examples of dysregulated epigenetic players include histone methyltransferases (e.g., SETD1B, SUV39H2, KMT2D, EZH2), histone demethylases (KDM6B) and chromatin remodeling factors (e.g., CHAF1A, ATAD2 CHAF1B), indicating that in addition to DNA methylation and histone acetylation (Ito et al. 2005; Szulakowski et al. 2006) other epigenetic layers may also be dysregulated in COPD.

### Integrative data analysis reveals epigenetically regulated genes in COPD fibroblasts

To further dissect the association between alterations in DNA methylation and changes in gene expression, we assigned DMRs to genes in their proximity. In total, we detected 4,059 genes associated with at least one DMR (in total 4,424 DMRs) within 4 kb upstream of the transcriptional start site (TSS) to 4 kb downstream of the transcriptional termination site (TTS) (**Figure 4A**). About 45% of the gene-associated DMRs are located close to the TSS, mainly in the promoter and first intron (**Figure 4B, Suppl. Fig. 4A-B**). To further decipher the interplay between expression and methylation changes in COPD, we focused our analysis on DMRs within 4 kb surrounding the TSS (**Figure 4C**). We observed an overrepresentation at differentially expressed genes compared to genes whose expression is not significantly changed in COPD (Fisher’s exact test: p-value = 1.3 x 10^-7^, **Suppl. Fig. 4C**). In total, 77 differentially expressed genes were associated with at least one DMR (**Figure 4C**), which was mainly located in the promoter or downstream of the TSS (**Suppl. Fig. 4B**). Spearman correlation between DNA methylation and expression change was calculated to assess how epigenetic variations are related to transcriptional differences. In contrast to unchanged genes which exhibited the expected normal distribution (**Figure 4D**, blue line), differentially expressed genes displayed a bimodal curve with enrichment at high positive and negative correlation rates, suggesting that some genes might be dysregulated by aberrant methylation in COPD (**Figure 4D**, red line). Examples of genes showing correlated DNA methylation and gene expression changes include UMPS, STEAP3, GABRR1, GLI4, AQP3, and LPXN (**Figure 4E**, **Suppl. Fig. 4D**), suggesting potential regulation of their expression by DNA methylation.

**Figure 4.**
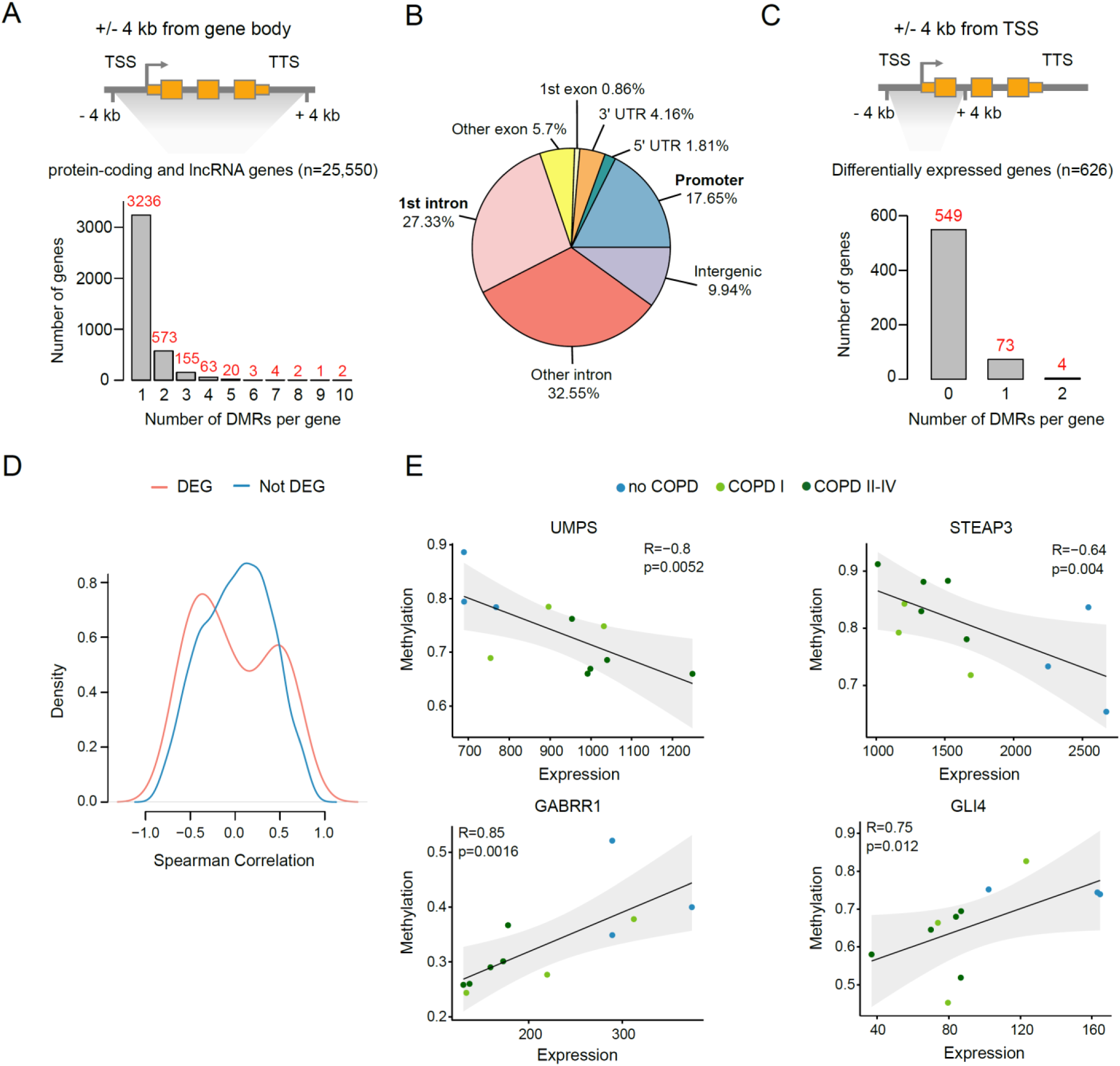
Integrative data analysis reveals epigenetically regulated genes in COPD fibroblasts. **A** DMRs located in the proximity of annotated protein-coding and lincRNA genes. DMRs within +/- 4 kb from gene body were assigned to their corresponding gene. TSS, transcription start site; TTS, transcription termination site. **B** Gene features of gene-associated DMRs. Promoter is defined as the region of - 1 kb to + 100 bp around the TSS. **C** DMRs located in the proximity of the TSS of DEGs. DMRs within +/- 4 kb from TSS of DEG were assigned to their corresponding gene. **D** Spearman correlation between gene expression and DMR methylation. DMRs within +/- 4 kb from TSS were considered. Gene-DMR pairs were split into DEGs (red) and not significantly changed genes (no DEG, blue). **E** Scatter plots showing examples of correlations between gene expression and methylation of promoter associated DMRs. Each dot represents an individual donor. Dots are color coded according to disease state. Gene expression is illustrated as normalized counts. Methylation is illustrated as average beta value of the corresponding DMR. Linear regression analysis was performed (black line) and the 95% confidence interval is indicated (grey area). P-value and Spearman correlation coefficient (R) are indicated.

### Functional siRNA screens identify novel regulators of COPD phenotypes in lung fibroblasts

Integration of DNA methylation and gene expression data, together with upstream regulator analysis using Ingenuity Pathway Analysis (Kramer et al. 2014) (https://digitalinsights.qiagen.com/products-overview/discovery-insights-portfolio/analysis-and-visualization/qiagen-ipa/) allowed us to select candidate regulators of the observed epigenetic and transcriptional changes in COPD HLFs. Overall, 110 candidates were manually selected for functional validation using phenotypic screens (**Suppl. Fig. 5A**, **Table 3**), 78% of them were not linked to COPD before. To determine the function of the selected candidates in key fibroblast processes related to COPD, high-content image-based phenotypic assays using small-interfering RNAs (siRNA)-mediated gene knockdown (KD) were carried out in primary human lung fibroblasts isolated from two normal healthy (NHLFs) and three COPD (DHLFs) donors (**Figure 5A**).

**Figure 5.**
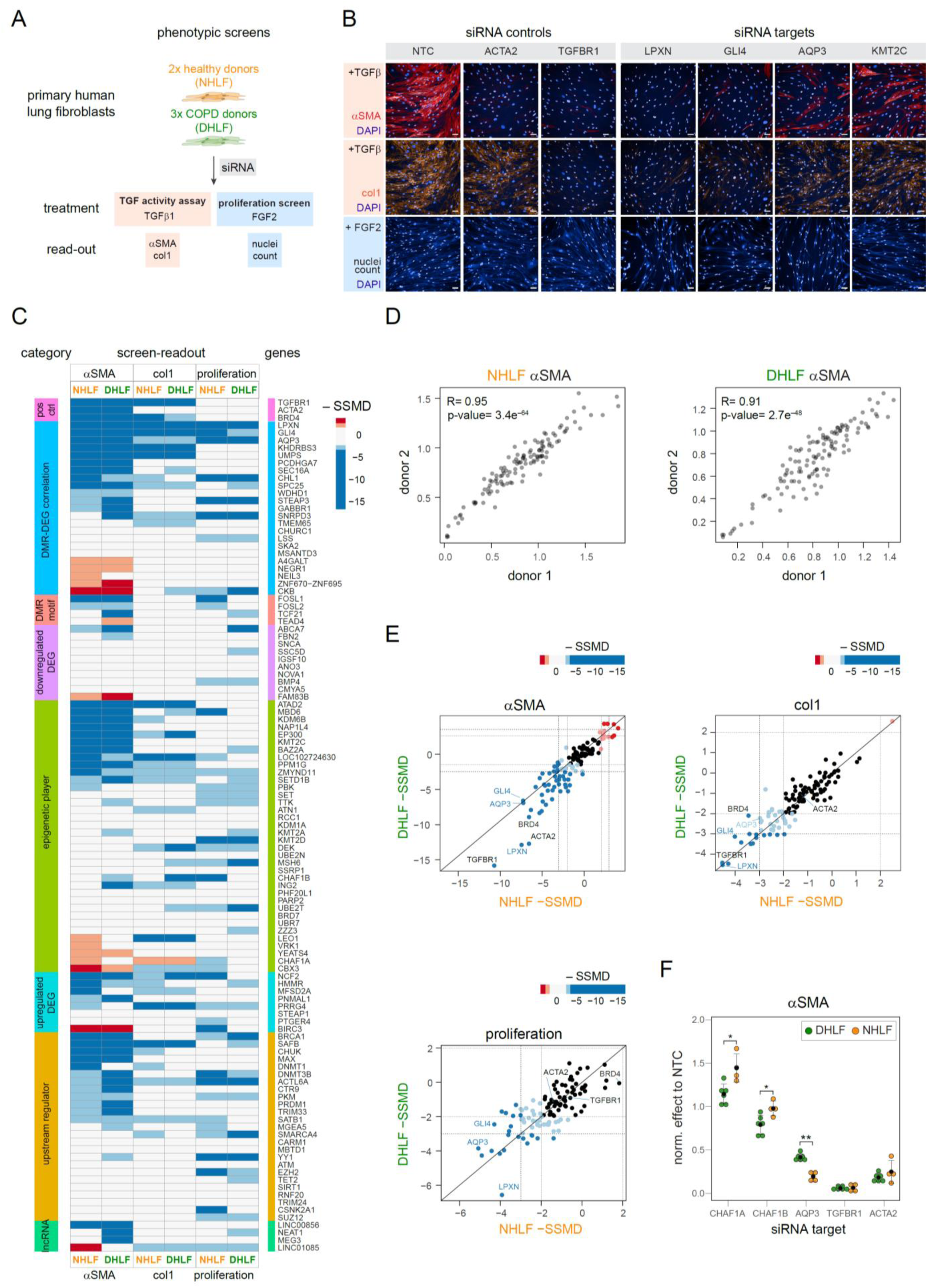
siRNA-based phenotypic screens in normal and COPD primary human lung fibroblasts identify multiple candidate genes regulating COPD phenotypes. **A** Schematic representation of the siRNA-based phenotypic assays performed in primary normal human lung fibroblasts (NHLFs, 2 donors) and diseased/COPD human lung fibroblasts (DHLFs, 3 donors). KD, knockdown; FGF, fibroblast growth factor; TGFβ, transforming growth factor beta; αSMA, alpha smooth-muscle actin; col1, collagen 1. **B** Examples of primary pictures obtained in the screens showing the performance of the siRNA controls as well as positive hits upon KD. NTC, non-targeting siRNA control. **C** Heatmap showing the effect of the KD of each candidate gene on the three measured readouts (αSMA, col1 and proliferation) in primary normal (NHLF) or COPD (DHLF) human lung fibroblasts, red: readout higher than NTC, blue: readout lower than NTC; SSMD = strictly standardized mean difference. |SSMD values| ≥ 2 are shown in lighter shade and |SSMD values| ≥ 3 in stronger shade. **D** Scatterplots showing the correlation of the screen data from 2 different NHLFs donors (left) and 2 different DHLFs (right) for the αSMA readout. **E** Comparison of the KD effect of each candidate relative to NTCs in NHLFs and DHLFs. Each dot represents a unique candidate tested, blue and red dots represent significant hits (|SSMD values| ≥ 2 are shown in lighter shade and |SSMD values| ≥ 3 in stronger shade). Assay controls are labeled in black and examples of strong hits regulating fibroblast to myofibroblast transition and cell proliferation processes are labeled in blue. **F** Dotplots showing examples of positive hits with significant differences between NHLFs (2 donors, 2 biological replicates each, shown in green) and DHLFs (3 donors, 2 biological replicates each, shown in orange) in αSMA readout. Screen readout was normalized to the corresponding non NTC. TGFβR1 and ACTA2 represent positive screen controls. The results of all replicates are shown. Statistical evaluation was performed with unpaired two-tailed student’s t-test. *: p-value < 0.05; **: p-value < 0.01. Black dots denote the means and error bars represent the standard deviation.

In COPD/emphysema, reduced proliferation, migration, and response to TGFβ1 of lung fibroblasts have been documented (Nobukuni et al. 2002; Holz et al. 2004; Muller et al. 2006; Togo et al. 2008), indicating impaired tissue-repair capacity of fibroblasts in the COPD lung. Reduced fibroblast activity in the injured alveolar microenvironment has been proposed as a critical mechanism driving the development of emphysema (Plantier et al. 2007; Kulkarni et al. 2016). Thus, to evaluate fibroblast response to TGFβ1 upon candidate gene knockdown, we performed a TGFβ1-induced fibroblast-to-myofibroblast transition assay (FMT). High-content image-based quantification of collagen 1 deposition (col1) and α-smooth-muscle fibers (αSMA), TGFβ1-responsive genes, were used as readouts. Additionally, to assess the proliferation capacity of primary fibroblasts upon candidate gene knockdown, we quantified the number of nuclei upon fibroblast growth factor 2 (FGF2) stimulation (**Figure 5A-B**).

The technical performance of the assays was demonstrated by the robust effect of the siRNAs targeting assay controls. As expected, siRNA targeting TGFβ receptor 1 (TGFβR1) showed a strong impact on the FMT assay (on both αSMA and col1 readouts) (**Figure 5B**). Similarly, siRNA against ACTA2 (which encodes αSMA) showed a specific effect only in the αSMA readout. Furthermore, the strong correlation between the results obtained from different donors among normal fibroblasts (e.g., R=0.95, p-value: 3.4×10^-64^ for αSMA) and fibroblasts derived from COPD patients (e.g., R=0.91, p- value: 2.7×10^-48^ for αSMA) confirmed the robustness of the assays (**Figure 5D** and **Suppl. Fig. 5B**).

To evaluate the differences between non-targeting siRNA controls (NTC) and gene-targeted knockdowns, we calculated strictly standardized mean differences (SSMD) (Zhang 2007; Zhang et al. 2007). Overall, a high hit-rate was observed, as 61 out of the selected 110 candidates showed an effect in at least one assay after applying the strict cutoff of |3 SSMD| and 87 when using a SSMD cutoff of |2|, demonstrating the power of multimodal analysis in identifying candidates with regulatory potential (**Figure 5C, 5E, Suppl. Fig. 5C, Table 4**). Three genes, leupaxin (LPX), aquaporin 3 (AQP3) and GLI4, showed strong effects in all three readouts in both normal and diseased cells, indicating their critical function in fibroblast biology (**Figure 5B-C and 5E**). Among the positive hits, multiple epigenetic factors were also present (**Figure 5C, Suppl. Fig. 5C, Table 4**). For example, we observed strong effects on lung fibroblast proliferation and differentiation upon targeted knockdown of different epigenetic enzymes (DNA methyltransferase DNMT3B, histone methyltransferases KMT2A, MKT2B, KMT3C, SETD1B, and EZH2, histone acetyltransferase EP300), chromatin remodeling factors (CHAF1A and CHAF1B) as well as epigenetics readers (BAZ2, CBX3), identifying these factors as key regulators of fibroblasts and COPD phenotypes. The imbalance of histone acetyltransferase (HAT) and deacetylase (HDAC) activities has previously been linked to COPD (Ito et al. 2005), providing a scientific basis for the potential use of bromodomain (BET) and HDAC inhibitors in COPD (van den Bosch et al. 2017), however the potential of targeting other dysregulated epigenetic activities in COPD remains to be explored.

Interestingly, we observed potential disease-specific effects for some of the tested candidates that were preserved between different donors. Here, the effect after siRNA-mediated gene knockdown varied between normal and diseased fibroblasts (**Figure 5F**, **Suppl. Fig. 5D**). For example, knockdown of CHAF1A increased, while knockdown of CHAF1B reduced the expression of αSMA, respectively, and both effects were stronger in diseased fibroblasts compared to normal cells. In turn, knockdown of AQP3 in normal fibroblasts had a larger effect on αSMA levels upon TGFβ1 stimulation compared to diseased cells (**Figure 5F**). All 3 genes were dysregulated in our RNA-seq in COPD (CHAF1A and CHAF1B were upregulated, whereas AQP3 was downregulated in COPD, **Table 2**), indicating that their dysregulation may be linked to COPD phenotypes in fibroblasts.

The cell-based assays in primary normal and COPD fibroblasts confirmed the functional role of numerous candidates identified from profiling data, indicating that integrating genome-wide epigenetic and transcriptomic profiling of purified normal and diseased human lung cells is a powerful approach for the identification of novel regulators of disease phenotypes. In addition, the presence of various epigenetic factors among the positive hits demonstrates that epigenetic regulation in COPD is an exciting research field that should be explored in-depth, as it holds promise for novel therapeutic avenues for patients with COPD.

## Discussion

In this study, we reveal novel transcriptomic and epigenetic signatures associated with COPD onset and progression, establishing a roadmap for further dissection of molecular mechanisms driving COPD phenotypes in lung fibroblasts.

Earlier studies using various patient material consisting of mixed-cell populations provided evidence of dysregulated DNA methylation patterns in COPD and identified CpG sites and pathways associated with smoking and COPD (Sood et al. 2010; Qiu et al. 2012; Vucic et al. 2014; Wan et al. 2015; Yoo et al. 2015; Busch et al. 2016; Morrow et al. 2016; Sundar et al. 2017; Carmona et al. 2018). Two recent publications also suggested that DNA methylation changes may originate in early life (Kachroo et al. 2020) and be linked to the severity of airflow limitation (Casas-Recasens et al. 2021). However, they all used low-resolution approaches, covering a representation of the genome only, mostly gene promoters. Hence, the full epigenomic landscape of COPD cells remains uncharted. Overall, there has been a limited consistency between different studies, likely coming from the cellular heterogeneity of the starting material, diverse donor selection criteria and different statistical models used. The dissection of cell-type specific mechanisms associated with COPD requires epigenetic profiling of defined cell populations. Only one study investigated DNA methylation changes in COPD patients with cell-type resolution (Clifford et al. 2018). Using Illumina 450K BeadChip Array (focusing on gene promoters), Clifford *et al*. identified 887 and 44 differentially methylated CpG sites in parenchymal and airway fibroblasts of COPD patients, respectively (Clifford et al. 2018). Our study, providing a much higher resolution of previously unexplored regions (e.g., enhancers) significantly extends these observations and demonstrates pronounced, genome-wide DNA methylation and gene expression changes in parenchymal fibroblasts in COPD, in both mild and severe disease.

Little is known about the correlation of DNA methylation with disease severity. Methylation changes in 13 genes have been identified in the lung tissue of COPD GOLD I and II patients compared to non-smoker controls (Casas-Recasens et al. 2021). However, it is unclear whether they represent smoking- or COPD-related changes, as ex-smoker controls were not investigated in this study (Casas-Recasens et al. 2021). Our data reveal that genome-wide DNA methylation changes are present in lung fibroblasts of COPD (I) patients compared to controls with matched smoking status and history (all ex-smokers), demonstrating that epigenetic changes occur early in disease development. Notably, COPD (I) samples clustered with COPD (II-IV) rather than no COPD samples, indicating that DNA methylation may provide a sensitive biomarker for early disease detection. This hypothesis awaits further validation in larger patient cohorts.

Currently, it is unclear how altered DNA methylation patterns in COPD translate into biological effects in COPD fibroblasts. DNA methylation in regulatory regions can modulate the binding of transcriptional factors to DNA (Stadler et al. 2011), hence, methylation profiling allows identifying transcriptional regulators potentially mediating the epigenetic alterations observed. We detected a significant enrichment of binding sites for TCF21 and FOSL2/FRA2 transcription factors in the DMRs overlapping with strong enhancers in COPD. TCF21 is a mesenchyme-specific basic helix-loop-helix transcription factor regulating multiple processes, including proliferation, extracellular matrix assembly, as well as secretion of pro-inflammatory mediators (Akama and Chun 2018). It is required for lung development in mice, mesenchymal-epithelial crosstalk (Quaggin et al. 1999) and specification of fibroblast cell fate in different organs (Acharya et al. 2012; Braitsch et al. 2012). Recently, TCF21 has been identified as a specific marker of lipofibroblasts in mouse (Park et al. 2019) and human lung (Liu et al. 2021), a subpopulation of fibroblasts essential for alveolar niche homeostasis and repair. Despite its central roles in alveolar development and maintenance, TCF21 function in human lung fibroblasts is largely unknown. Our phenotypic screens demonstrate that TCF21 is required for lung fibroblast proliferation and differentiation upon TGFβ1 stimulation, providing first insights into its molecular function. Notably, the effects of TCF21 on proliferation and αSMA were stronger in diseased cells, indicating that COPD cells may be more sensitive to TCF21 loss that healthy lung fibroblasts. Consistent with the documented role of TCF21 in regulating cell proliferation in cancer (Lotfi et al. 2021), its differential binding at strong enhancers in COPD fibroblasts could provide a molecular mechanism for the differential expression of DNA replication genes observed in our RNA-seq analysis.

FOSL2/FRA2 belongs to the activator-protein (AP)-1 family of transcription factors and is a known regulator of wound repair, and TGFβ mediated fibrosis (Eferl et al. 2008). Increased FOSL2/FRA2 expression is detected in several chronic lung diseases, including pulmonary fibrosis, COPD, and asthma (Birnhuber et al. 2019). Notably, ectopic expression of FOSL2/FRA2 in mice results in fibrosis of several organs, including the lung, highlighting a potential profibrotic role of FOSL2/FRA2 (Eferl et al. 2008). Furthermore, our high-content screens demonstrate that FOSL2/FRA2 is required for myofibroblast differentiation, consistent with its postulated profibrotic role. Collectively, our data suggest that TCF21 and FOSL2/FRA2, whose binding sites are enriched in DMRs at strong enhancers, may mediate downstream biological effects in COPD fibroblasts and contribute to disease phenotypes, linking epigenetic changes to gene regulatory networks.

Transient TGFβ1 activity is required for lung tissue regeneration and repair upon injury, however its persistent activation in lung fibroblasts leads to aberrant repair and fibrosis (Fernandez and Eickelberg 2012). In turn, reduced fibroblast proliferation and response to TGFβ1 has been proposed as a key mechanism driving the development of emphysema (Plantier et al. 2007; Togo et al. 2008; Konigshoff et al. 2009; Kulkarni et al. 2016). How these phenotypes are controlled at the molecular level is not well understood. We identified and functionally validated numerous, previously unknown regulators of lung fibroblast function in COPD. Among the top candidates, silencing of the water/glycerol channel aquaporin 3 (AQP3), Hedgehog transcription factor GLI4 and focal adhesion protein leupaxin (LPXN) had the most drastic effects on both fibroblast proliferation and TGFβ1−mediated differentiation, establishing these three proteins as new regulators of lung fibroblast repair and remodeling. AQP3 contribution to wound healing, ECM remodeling and cell proliferation has been well documented in other cellular systems (Xu et al. 2011; Ryu et al. 2012; Chen et al. 2014; Huang et al. 2015; Hou et al. 2016; Luo et al. 2016; Xiong et al. 2017) and the role of LPXN in cancer cell proliferation and migration through regulation of focal adhesion sites is recognized (Kaulfuss et al. 2008; Dierks et al. 2015). Hence, downregulation of AQP3 in COPD fibroblasts could contribute, at least in part, to the decreased proliferation and contractility, manifesting in reduced fibroblast activity and impaired response to injury in emphysema (Togo et al. 2008). In support of our data, change in expression of AQP3 (Heinbockel et al. 2018) and LPXN (Spira et al. 2004) in COPD/emphysematous lung tissue has been observed previously, consistent with their dysregulation in COPD fibroblasts in our RNA-seq. Notably, we showed that changes in expression of AQP3, LPXN and GLI4 in COPD fibroblasts were also associated with aberrant methylation in proximity to their TSS, indicating that their epigenetic regulation may be one of the factors contributing to the reduced repair capacity of lung fibroblasts in emphysema (Muller et al. 2006; Togo et al. 2008).

Collectively, our results demonstrate that focused, high-resolution profiling of defined cell populations in COPD effectively complements large-cohort epigenetic biomarker studies and can provide important insights into COPD-driving cell populations and associated mechanisms. Integration of - omics data across disease stages is a powerful tool for the identification of novel candidate disease regulators and sensitive biomarkers. Future large-scale profiling of early disease is crucial for an improved understanding of COPD pathology and will guide the development of new diagnostic strategies and disease-modifying therapies.

## Data availability

The WGBS and RNA-seq data generated in this study have been deposited at the European Genome-phenome Archive (EGA), which is hosted by the EBI and the CRG, under accession EGAXXXXXXX.

## Methods

### Study approval

The protocol for tissue collection was approved by the ethics committees of the University of Heidelberg (S-270/2001) and Ludwig-Maximilians-Universität München (projects 333-10 and 17-166)

the University of Texas Health Science Center at Houston (HSC-MS-08-0354 and HSC-MS-15-1049) and followed the guidelines of the Declaration of Helsinki. All patients gave written informed consent before inclusion in the study and remained anonymous in the context of this study.

### Patient samples

Lung tissue samples were obtained through collaborations with the Lung Biobank Heidelberg at Thorax Clinic (Heidelberg, Germany), the Asklepios Clinic (Gauting, Germany) and the UTHealth Pulmonary Center of Excellence (Houston, TX, USA). Residual lung parenchyma samples were obtained from patients undergoing lung surgery due to primary squamous cell carcinomas (SCC) who had not received chemotherapy or radiation within 4 years before surgery or from COPD patients undergoing lung resection. Normal human lung tissue used for protocol optimization was obtained from the International Institute for the Advancement of Medicine (IIAM), from lungs rejected for transplantation due to reasons unrelated to obvious acute or chronic pulmonary disease.

### Collection of lung tissue samples for profiling

To identify molecular changes associated with COPD development and progression we collected distal lung tissue from patients across COPD stages and divided them into three groups: 1) no COPD, 2) mild COPD (stage I, according to GOLD classification(2021)) and 3) established COPD (GOLD stages II-IV, **Suppl. Fig. 1A**). Strict patient inclusion criteria for a prospective tissue collection were established to ensure best possible matching of control and disease groups. To avoid direct smoking effects (van der Vaart et al. 2004), all included donors were ex-smokers. In addition, lung function results, as well quantitative emphysema score index (ESI) based on chest CT and whenever possible, medical history were collected for each patient for their best possible characterization. Patients’ characteristics and representative images from hematoxylin and eosin staining of the tissue are provided in **Suppl. Fig. 1A and 1B**, respectively. Each tissue sample was reviewed by an experienced lung pathologist, who confirmed that all samples were tumor-free and evaluated COPD relevant phenotypes, like emphysema, airway thickening and immune infiltration. Only ex-smokers with preserved lung function and no indication of emphysema or fibrosis in the test results or patient history were included as control samples. Importantly, we included 2 samples from COPD (GOLD II-IV) donors (HLD38 and HLD39), which originated from lung resections, ensuring that the observed changes are present in COPD tissue without cancer background.

There were no significant differences between control and COPD donors regarding age, body mass index, smoking status, and smoking history, but the control and COPD group could be clearly separated based on lung function data (**Figure 1A** and **Suppl. Fig. 1A**). Tissue samples that met the inclusion criteria were cryopreserved upon collection to allow their thorough characterization by an experienced lung pathologist before cell isolation and profiling (**Figure 1B** and **Suppl. Fig. 1B**). We have previously shown that this step is crucial to ensure exclusion of low-quality control samples presenting additional lung pathologies, which may result in confounding effects in sequencing-based analyse (Llamazares-Prada et al. 2021).

### Emphysema score index (ESI) determination

Lung and emphysema segmentation were performed to calculate the ESI from clinically indicated preoperative CT scans taken with mixed technical parameters. After automated lung segmentation using the YACTA-software, a threshold of -950 HU was used with a noise-correction range between - 910 and-950 HU to calculate the relative amount of emphysema in % of the respective lung portion (Lim et al. 2016). While usually global ESI was measured, only the contralateral non-affected lung side was used if one lung was severely affected by the tumor.

### FFPE and H&E

Representative slices from different areas of the tissue were fixed O/N with 10% neutral buffered formalin (Sigma-Aldrich). Next, fixed tissue samples were washed with PBS (Fisher Scientific) and kept in 70% ethanol at 4°C. Sample dehydration, paraffin embedding, and hematoxylin and eosin (H&E) staining was performed at Morphisto (Morphisto GmbH, Frankfurt, Germany). Per sample, two 4 µm thick sections were cut on a Leica RM2255 microtome with an integrated cooling station and water basin and transferred to adhesive glass slides (Superfrost Plus, Thermo Fisher). Subsequently, the sections were dried O/N in a 40°C oven to remove excess water and enhance adhesion. H&E- stained slides were evaluated by an experienced lung pathologist at the Thorax Clinic in Heidelberg.

### Cryopreservation of lung parenchyma

Lung tissue was cryopreserved upon reception as described previously (Llamazares-Prada et al. 2021). Briefly, specimens were transported in CO_2_-idepenent medium (Thermo Fisher Scientific) supplemented with 1% BSA (Carl Roth), 1% penicillin & streptomycin (Fisher Scientific) and 1% Amphotericin B (Fisher Scientific). Upon reception, tissue pieces were carefully inflated with ice-cold HBSS (Fisher Scientific), supplemented with 2mM EDTA (Thermo Fisher Scientific), 1% BSA (Carl Roth), 1% penicillin & streptomycin (Fisher Scientific) and 1% Amphotericin B (Fisher Scientific). Exemplary samples of the different areas of the lung piece were collected for subsequent histological analysis. The pleura was removed from the remaining tissue, and airways and vessels separated from the parenchyma as much as possible. The parenchymal airway and vessel-free fractions were minced, transferred to cryo-tubes, covered with ice-cold freezing medium [70% DMEM, high glucose with GlutaMAX^TM^ (Thermo Fisher Scientific), 20% FBS (Gibco) and 10% DMSO (Carl Roth)], kept on ice for 15 min, and transferred to -80°C in Mr. Frosty^TM^ containers (Nalgene) to ensure a gradual temperature decrease (1°C/min). For long-term storage, samples were kept in liquid nitrogen.

### Fibroblast isolation from human lung tissue

As no universal fibroblast markers for FACS are available, primary human lung fibroblasts were isolated by explant outgrowth from tumor-free, distal parenchymal lung tissue that has been depleted from visible airways and vessels, as previously described (Llamazares-Prada et al. 2021). Briefly, 7-8 micro-dissected lung parenchyma pieces were placed per well into 6-well plates, let for 30 min at RT without medium to improve explant attachment, and carefully covered with 1 mL of growth medium: DMEM, high glucose, GlutaMAX^TM^ (Thermo Fisher Scientific) supplemented with 2% FBS (Gibco) and 1% penicillin & streptomycin (Thermo Fisher Scientific). Explants were left undisturbed for 4-7 days, afterwards the medium was exchanged every 2 days and the outgrowth of fibroblasts from the explants was followed daily. Cells were collected from multiple explant pieces when reaching 70% confluency to preserve the fibroblast heterogeneity. Possible epithelial contamination was prevented by short trypsinization (0.05% trypsin with EDTA (Gibco), 3 min at 37°C) and keeping cells in the growth medium indicated above, suitable for fibroblasts enrichment.

### Immunofluorescence of human lung fibroblasts

The purity of the isolated fibroblasts was assessed by immunofluorescence using mesenchymal markers vimentin (VIM) and alpha smooth muscle actin (αSMA) as follows. 10^4^ human lung fibroblasts in passage 3 were seeded per well in a 96-well plate for imaging (Zell-kontakt). 48h later, cells were washed with 1X PBS (Fisher Scientific), fixed for 10 min with 4% PFA (Sigma-Aldrich) at RT, washed and permeabilized for 10 min with 0.3% Triton-X-100 (Carl Roth) at RT. Unspecific staining was blocked by incubating 1h at RT with blocking buffer: 5% BSA (Carl Roth), 2% Normal Donkey Serum (Abcam) in 1X PBS (Fisher Scientific). Cells were incubated with primary antibodies against VIM (sc- 7557, Santa Cruz Biotechnology, 1:200) and αSMA (ab7817, Abcam, 1:100) overnight at 4°C and labelled with respective secondary antibodies [donkey anti-goat IgG Alexa Fluor 488 (A-11055, Thermo Fisher Scientific, 1:500) and donkey anti-mouse IgG Alexa Fluor 568 (A-10037, Thermo Fisher Scientific, 1:500)] for 40 min at RT in the dark. After washing with 1X PBS (Fisher Scientific), the nuclei were counterstained with DAPI (Thermo Fisher Scientific, 1:5,000) for 10 min at RT and washed with 1X PBS. Stained and fixed cells were kept in 1X PBS (Fisher Scientific) at 4°C in the dark until imaging. Imaging was conducted at the ZMBH imaging facility (Heidelberg, Germany) using the Zeiss LSM780 confocal fluorescent microscope.

### FACS analysis of isolated fibroblasts and lung suspension

Cryopreserved lung tissues were thawed for 2 min in a 37°C water-bath, collected in 50mL Falcon tubes and washed with wash buffer: HBSS supplemented with 2mM EDTA (Thermo Fisher Scientific), 1% BSA (Carl Roth), 1% penicillin & streptomycin (Fisher Scientific) and 1% Amphotericin B (Fisher Scientific). Tissue was minced into smaller pieces prior to mechanical and enzymatic dissociation as indicated previously(Llamazares-Prada et al. 2021). Briefly, the minced tissue (1g) was introduced in GentleMACS C-tubes (Miltenyi Biotec) containing 10 μM ROCK inhibitor (Y-27632, Adooq Bioscience), 10 μg DNAseI (ProSpec-Tany TechnoGene), the enzyme mix from the human tumor tissue dissociation kit (Miltenyi Biotec) and 4.5 mL of CO_2_-independent media (Thermo Fisher Scientific) supplemented with 1% BSA (Carl Roth), 1% penicillin & streptomycin (Fisher Scientific) and 1% Amphotericin B (Fisher Scientific). Tubes were closed tightly, introduced into the GentleMACS dissociator (Miltenyi Biotec) for mechanic disruption and the following program was performed: program h_tumor_01, followed by 15 min incubation at 37°C on a rotator; h_tumor_01, 15 min at 37°C on a rotator; h_tumor_02, and 15 min at 37°C on a rotator for a final enzymatic dissociation and a last mechanical shearing using the program h_tumor_02. The samples were pipetted up and down to help disaggregating. Finally, the enzymatic reaction was stopped by adding 20% FBS (Gibco) and single cells were collected by sequential filtering through 100 µm, 70 µm, and 40 µm cell strainers (BD Falcon). Cells were centrifuged, resuspended in ACK lysis buffer (Sigma Aldrich) and incubated for 3 min at RT to lyse erythrocytes. Lung single-cell suspensions were washed with HBSS (Fisher Scientific) supplemented with 2mM EDTA (Thermo Fisher Scientific), 1% BSA (Carl Roth), 1% penicillin & streptomycin (Fisher Scientific) and 1% Amphotericin B (Fisher Scientific).

To generate fibroblast single-cell suspensions, passage 3 fibroblasts were trypsinized 5 min at 37°C using 0.05% trypsin with EDTA (Gibco), centrifuged at 1000 rpm for 5 min at RT and resuspended in HBSS (Fisher Scientific) supplemented with 1% BSA (Carl Roth), 1% penicillin & streptomycin (Fisher Scientific) and 1% Amphotericin B (Fisher Scientific).

Lung and fibroblast single-cell suspensions were incubated with human TruStain FcX (Biolegend) for 30 min on ice to block Fc receptors. Immune and epithelial cells were labelled using CD45 (CD45- Bv605, BD Bioscience) and EpCAM (anti-human CD326 -PE, Affymetrix eBioscience) antibodies respectively for 30 min in the dark at 4°C following manufacturer instructions. Stained samples were washed with PBS 1X (Fisher Scientific) and resuspended in HBSS (Fisher Scientific) supplemented with 2mM EDTA (Thermo Fisher Scientific), 1% BSA (Carl Roth), 1% penicillin & streptomycin (Fisher Scientific) and 1% Amphotericin B (Fisher Scientific). Stained cells were added to Falcon 5 mL polystyrene tubes with 40 µm cell strainer caps (Neolab Migge). To discriminate between live and dead cells, we used SyTOX blue (Thermo Fisher Scientific) as recommended by manufacturer.

### RNA isolation and RNA-seq

10^5^ HLFs were harvested at passage 3 for RNA-seq studies by scraping in ice-cold PBS. Total RNA was isolated using RNeasy plus micro kit (Qiagen, Hilden, Germany) following manufacturer’s instructions. DNA was removed by passing the lysate through the gDNA eliminator column and by an additional on-column DNase I treatment (Qiagen) before the elution. RNA was eluted using nuclease- free water (Thermo Fisher Scientific) and the concentration measured with Qubit HS Kit (Thermo Fisher Scientific). RNA integrity was assessed using the Bioanalyzer 2100 (Agilent, model G2939A) and the RNA 6000 pico kit (Agilent). Only samples with RIN > 8.5 were processed.

Libraries were prepared at the Genomics core facility (GeneCore) at EMBL using 200 ng of total RNA as input. Ribosomal RNA was removed by Illumina Ribo-Zero Gold rRNA Removal Kit Human/Mouse/Rat (Illumina, San Diego, CA, USA) and stranded total RNA-seq libraries were prepared using the Illumina TruSeq RNA Sample Preparation v2 Kit (Illumina, San Diego, CA, USA) implemented on the liquid handling robot Beckman FXP2. Obtained libraries were pooled in equimolar amounts. 1.8 pM solution of each library was pooled and loaded on the Illumina sequencer NextSeq 500 High output and sequenced uni-directionally, generating ∼450 million reads per run, each 75 bases long.

### RNA-seq read alignment and transcript abundance quantification

Single-end reads were mapped to the human genome version 37 (hg19) and the reference gene annotation (release 70, Ensembl) using STAR v2.5.0a (Dobin et al. 2013) with following parameters:

--outFilterType BySJout --outFilterMultimapNmax 20 --alignSJoverhangMin 8 -- alignSJDBoverhangMin 1 --outFilterMismatchNmax 999 --alignIntronMin 20 --alignIntronMax 100000 - -outFilterMismatchNoverReadLmax 0.04 --outSAMtype BAM SortedByCoordinate --outSAMmultNmax 1 --outMultimapperOrder Random.

Contamination of PCR duplication artefacts in the RNA-seq data was controlled using the R package dupRadar (Sayols et al. 2016). The *featureCounts* script (Liao et al. 2014) of the Subread package v1.5.3 was used to assign and count mapped reads to annotated protein-coding and lncRNA genes with default settings.

### Differential Gene Expression Analysis

Statistical analysis of differential gene expression was performed with the DESeq2 Bioconductor package (Love et al. 2014). For exploratory RNA-seq data analysis the data needs to be homoscedastic. Therefore, the raw counts were transformed by the regularized-logarithm transformation *rlog*. Genes with less than 32 counts in at least 5 samples were excluded from further analysis. Ex-smokers with preserved lung function (no COPD) were considered as ground state and differential gene expression in COPD patients classified as GOLD Grade II-IV was identified as a significant change in expression by an FDR (false discovery rate) < 0.05 and an absolute log2 fold change > 0.5 (corresponding to fold change > 1.4), after fold change correction with the built-in *lfcShrink* function.

### DNA isolation and T-WGBS

Genomic DNA was extracted from 2×10^5^ primary human lung fibroblasts harvested in passage 3 using QIAamp Micro Kit (Qiagen, Hilden, Germany) following manufacturer’s protocol, with an additional RNase A treatment step. T-WGBS was essentially performed as described previously (Wang et al. 2013) using 30 ng genomic DNA as input. 15 pg unmethylated DNA of phage lambda was used as control for bisulfite conversion. Four sequencing libraries were generated per sample using 11 amplification cycles. For each sample, equimolar amounts of all four libraries were pooled and sequenced on two lanes of a HiSeq2500 (Illumina, San Diego, California, US) machine at NGX Bio (San Francisco), resulting in 100 bp, paired-end reads.

### Read Alignment

The whole genome bisulfite sequencing mapping pipeline MethylCtools with modifications to adapt for the T-WGBS data was used (https://github.com/hovestadt/methylCtools) (Hovestadt et al. 2014). Briefly, the hg19 reference genome (37d5) was transformed *in silico* for both the top strand (C to T) and bottom strand (G to A). Before alignment, adaptor sequences were trimmed using Trimmomatic (release 0.35) (Bolger et al. 2014). The first read in each read pair was then C-to-T converted and the 2nd read in the pair was G-to-A converted. The converted reads were aligned to a combined reference of the transformed top (C to T) and bottom (G to A) strands using BWA MEM (bwa-0.7.8) with default parameters, yet, disabling the quality threshold for read output (-T 0) (Li and Durbin 2009). After alignment, reads were converted back to the original states, and reads mapped to the antisense strand of the respective reference were removed. Duplicate reads were marked, and the complexity determined using Picard *MarkDuplicates* (http://picard.sourceforge.net/). Total genome coverage was calculated using the total number of bases aligned from uniquely mapped reads over the total number of mappable bases in the genome.

### Methylation calling

At each cytosine position, reads that maintain the cytosine status were considered methylated, and the reads that have cytosine converted to thymine were considered unmethylated. Only bases with Phred-scaled quality score of ≥ 20 were considered. In addition, the 10 bp at the two ends of the reads were excluded from methylation calling according to M-bias plot quality control. In addition, CpGs located on sex-chromosomes were removed from analysis.

### DMR calling

Differences in CpG methylation profiles of no COPD donors and patients diagnosed with COPD were analyzed using the R/Bioconductor package bsseq (Hansen et al. 2012). First, the data was smoothed using the built-in *BSmooth* function with default settings. Only CpG sites with a coverage of at least 4x were kept for subsequent analysis. A t-statistic was calculated between no COPD and COPD (II-IV) samples using the *BSmooth.tstat* function with following parameters: local.correct=TRUE, maxGap=300, estimate.var=”same”. Differentially methylated regions (DMRs) were called by 1) selecting the regions with the 5% most extreme t-statistics in the data (lower and upper 2.5% quantile; default paramters of the dmrFinder function), 2) filtering for regions exhibiting at least 10% methylation difference between no COPD and COPD (II-IV) and containing at least 3 CpGs. 3) Finally, a non-parametric Wilcoxon test was applied using the average methylation level of the region to remove potentially false positive regions, since the t-statistic is not well-suited for not normally distributed values, as expected at very low/high (close to 0% / 100%) methylation levels. A significance level of 0.1 was used.

### Gene ontology analysis

The closest genes were assigned to DMRs and subjected to gene ontology enrichment analysis using GREAT (McLean et al. 2010)

### ChIP-seq Data

Histone modification ChIP-seq data of human lung fibroblasts were obtained from the ENCODE portal (https://www.encodeproject.org/) (Davis et al. 2018) with the following identifiers: ENCFF354IJB, ENCFF070CZY, ENCFF377BNX, ENCFF227WSF, ENCFF208SHP, ENCFF386FDQ, ENCFF102BGI, ENCFF843AYT.

Chromatin states for adult human lung fibroblasts (accession: ENCFF001TDQ) were obtained from ENCODE data base (Ernst and Kellis 2017). The number of bp of each DMR coinciding with a chromatin state was calculated and the chromatin state with the largest overlap was assigned to the DMR. To assess the genomic background, 10.000 regions with matching size and CpG distribution were randomly selected.

### Profile Plots of DMRs and Identification of Super-enhancers

Enrichment of H3K4me1, H3K27ac and H3K27me3 signals at DMRs, stratified in hypo- and hypermethylated regions, was performed with peakSeason (https://github.com/PoisonAlien/peakseason).

Super Enhancers (SE) were identified using ROSE (Rank Ordering of Super Enhancers) software (v.0.1; https://bitbucket.org/young_computation/rose), by merging closely spaced (<12.5 kb) enhancer peaks (H3K4me1 peaks overlapping with H3K27ac peaks) (Whyte et al. 2013). Further on, all enhancers were ranked by their H3K27ac signals. Separation of SE and enhancers was performed based on the geometrical inflection point.

### Transcription factor motif analysis

All hypomethylated DMRs which showed an overlap with the strong enhancer chromatin state were selected and motif enrichment analysis was carried out using the *findMotifsGenome.pl* script of the HOMER software suit omitting CG correction.

In order to obtain information about methylation dependent binding for transcription factor motifs which are enriched at DMRs, the results of a recent SELEX study (Yin et al. 2017) were integrated in the analysis and a motif database of 1787 binding motifs with associated methylation dependency was constructed. The log odds detection threshold was calculated for the HOMER motif search as following:

Bases with a probability > 0.7 get a score of log(base probability/0.25), otherwise the score was set to 0. The final threshold was calculated as the sum of the scores of all bases in the motif. Motif enrichment analysis was carried out against a sampled background of 50,000 random regions with matching GC content using the *findMotifsGenome.pl* script of the HOMER software suit omitting CG correction and setting the generated SELEX motifs as motif database.

### siRNA-based phenotypic assays in primary human lung fibroblasts

Normal human lung fibroblasts (NHLFs) from 2 donors (donor IDs: 608197; 543644) and diseased COPD human lung fibroblasts (DHLFs) from 3 donors (donor IDs: OF3353, OF3418 and OF3238) were purchased from Lonza and tested for their response to FGF2 and TGFβ stimulation.

### Fibroblast to myofibroblast transition (FMT) assay

Cells in passage 5 were plated in a poly-D-lysine coated 384 CellCarrier microtiter plate from PerkinElmer in fibroblast basal medium (FBM) with FGM-2TM Single Quots (Lonza) at a density of 2000 cells per well. Six hours after cell seeding, cells were transfected with siRNAs (Horizon ON- Target Plus siRNA pools) as previously described (Weigle et al. 2019). 24h later, the medium was replaced by FBM containing 0,1% fetal calf serum (starvation medium). 24h later, fibroblast to myofibroblast differentiation was initiated by adding fresh starvation medium containing a mixture of Ficoll 70 and 400 (GE Healthcare; 37,5mg/mL and 25mg/mL, respectively), 200µM vitamin C and 5ng/mL TGFβ1. After 72h the medium was removed, cells were fixed with 100% ice-cold methanol for 30 min, washed with PBS, permeabilized 20 min using 1% Triton-X-100 (Sigma), washed, and blocked for 30 min with 3% BSA in PBS. After an additional wash step, cell nuclei were stained using 1µM Hoechst 33342 (Molecular Probes). Alpha smooth-muscle actin (αSMA) and collagen I (col1) were stained using monoclonal antibodies (1:1000 diluted, Sigma, A2547 and SAB4200678, respectively). For detection of primary antibodies, cells were washed and incubated for 30 min at 37°C with AF647-goat-anti-mouse IgG2b (αSMA) and AF568 goat-anti-mouse IgG1 (col1) antibodies. After removal of secondary antibodies, cells were stained with HCS Cell Mask Green stain (Invitrogen, 1:50,000). Following a final PBS 1x wash step, images were acquired in a GE Healthcare InCell 2200 Analyzer, using 2D-deconvolution for nuclei (Hoechst channel), cells (FITC channel), αSMA (Cy5 channel) and collagen I (TexasRed channel), and images were transferred to and analyzed using Perkin Elmeŕs Columbus^TM^ Image Storage as previously described(Aumiller et al. 2017; Weigle et al. 2019). Briefly, the building blocks (BB) of the Columbus^TM^ Image Analysis system were used, first nuclei (Hoechst channel acquisition) were detected using the BB “nuclei”. Second, cells were defined with the BB “find cytoplasm” from the FITC channel image. αSMA fibers and col1 area were defined by two individual BBs “find simple image region” based on images acquired in the Cy5 and TexasRed channels, respectively. Both αSMA fibers and col1 readouts were normalized to the number of cells per image field. The FMT assay was performed in 2 NHLFs and 3 DHLFs independent donors. For each donor, siRNA transfection for every gene was performed in 4 technical replicates. In addition, the FMT screen was performed in each donor twice independently.

### Proliferation assay (nuclei count)

To analyze the effects of gene knockdown on FGF2-mediated fibroblast proliferation, 2000 cells at passage 5 were plated in a poly-D-lysine coated 384-CellCarrier microtiter plate (PerkinElmer) in FBM with FGM-2TM Single Quots (Lonza). Six hours after seeding, cells were transfected with siRNAs (Horizon ON-Target Plus siRNA pools) as previously described(Weigle et al. 2019). 24h later, the medium was replaced by FBM containing 0,1% fetal calf serum (starvation medium). 24h later the medium was replaced by starvation medium containing 20ng/mL basic FGF (R&D Systems). After 72h the medium was removed, cells washed with PBS and treated with 3.7% formaldehyde containing 1µM Hoechst 33342 for 30 min. Cells were washed with PBS and images were acquired in a GE Healthcare InCell 2200 Analyzer, using 2D-deconvolution for nuclei (Hoechst channel). Nuclei numbers were determined using the Columbus^TM^ image analysis software as described above (BB “nuclei”). The proliferation assay was performed in 5 independent donors: 2 NHLFs and 3 DHLFs. For each donor, siRNA transfection to knockdown the selected candidate genes was performed in 4 technical replicates. In addition, the proliferation screen was performed in each donor twice independently.

### Analysis of phenotypic screen data

siRNA transfection was performed in each phenotypic screen in 4 technical replicates and repeated 2 times independently for each donor. For statistical analysis of both the FMT and proliferation data, each readout (nuclei for both FMT and proliferation, αSMA and col1 for FMT) was first normalized within each plate, based on the negative control wells, corresponding to cells transfected with non-target siRNA control (NTC) (40 wells per plate). After plate-based normalization, the normalized values for the specific readout (e.g., nuclei, αSMA and col1) were averaged for the independent replicates. To measure the siRNA effect as the magnitude of the difference between an individual siRNA and the negative control (NTC siRNA), the previously described strictly standardized mean difference (SSMD) was applied(Zhang 2007; Zhang et al. 2007). The following formula was used forthe SSMD calculation: 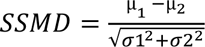, where µ1 is the normalized mean of all NTC siRNAs, µ2 is the mean of the normalized values of siRNA for a given gene, 𝜎1 is the variance of all normalized NTC siRNAs values and 𝜎2 is the variance of all normalized values transfected with siRNA for a given gene.

### Statistical analysis

An unpaired non-parametric *t* test (Mann-Whitney test, GraphPad Prism software, version 8.0.1) was employed to compare the lung function (FEV1 and FEV1/FVC values) between control and COPD donors (**Fig 1A**). For the **Suppl. Fig. 1A** displaying all the patient metadata of the three groups studied (control, COPD I and COPD II-IV), one-way ANOVA non-parametric unpaired test was used (Kruskal-Wallis test, GraphPad Prism software, version 8.0.1) followed by correction for multiple comparisons using Dunn’s test. The adjusted p-value for each comparison is shown. The significance level was set to 0.05.

## Author contributions

RZJ, US & MLP contributed to the design and conception of the study. MLP, SP and MR performed experiments with the help from VM, RT, MR, OS and DWe. MSchu and CK performed the phenotypic screens. US performed bioinformatic analysis and integration of the RNA-seq and WGBS data, with the help of KQ, JH and GP. TMu, MS, FH, HW, FH, HKQ and IK provided lung tissue and patient data. AW performed the pathological analysis of the H&E lung specimens. CPH determined the emphysema score index of the patients. HS, BJ, VB, DWy, TPJ, BM, BB, CI and CP provided critical input, analysis software and materials. RZJ, US and MLP wrote the manuscript with input from all authors. All authors contributed to scientific discussions and approved the final version of the manuscript.

## Acknowledgments

We would like to thank Lung Biobank (Heidelberg, Germany) – a member of the Biomaterial bank Heidelberg (BMBH), the tissue bank of the National Center for Tumor Diseases (NCT) and the Biobank platform of the German Center for Lung Research (DZL), as well as the Asklepios Biobank for Lung Diseases, member of the German Center for Lung Research (DZL) for providing Biomaterials and Data. We also thank Christa Stolp for help with collecting primary material. We acknowledge excellent sequencing service and helpful discussions from the Genomics core facility (GeneCore, EMBL, Germany) for RNA-seq and from NGX Bio (San Francisco, USA) for T-WGBS sequencing, as well as support from the ZMBH imaging facility (Heidelberg, Germany) for immunofluorescence. We thank Morphisto GmbH (Frankfurt, Germany) for excellent histological service. We also thank Christian Tidona (BioMed X Innovation Center) and Markus Koester (Boehringer Ingelheim) for helpful project discussions. We thank the ENCODE Consortium for providing and the Bradly Bernstein Lab for producing the ChIP datasets of human fibroblasts used in this study.

## Potential Conflict of Interest

RZJ, MLP, VM, US, RT, MR, SP, OS and BM as employees of BioMed X Institute received research funding by Boehringer Ingelheim Pharma GmbH & Co KG. HS, BJ, MSchu, CK, KQ and DWy are employees of Boehringer Ingelheim Pharma GmbH & Co KG and receive compensation as such. TM received a research grant, non-financial support and has patent applications with Roche Diagnostics GmbH outside of the described work. CPH has stock ownership in GSK; received research funding from Siemens, Pfizer, MeVis and Boehringer Ingelheim; consultation fees from Schering-Plough, Pfizer, Basilea, Boehringer Ingelheim, Novartis, Roche, Astellas, Gilead, MSD, Lilly Intermune and Fresenius, and speaker fees from Gilead, Essex, Schering-Plough, AstraZeneca, Lilly, Roche, MSD, Pfizer, Bracco, MEDA Pharma, Intermune, Chiesi, Siemens, Covidien, Boehringer Ingelheim, Grifols, Novartis, Basilea and Bayer, outside the submitted work. HW received consultation fees from Intuitive and Roche.

## Funding

This study was supported by Boehringer Ingelheim. The work was partly funded by the School of Biosciences (Cardiff University) to RZJ and the German Center for Lung Research (DZL) to CP, TM, MS, FH, CPH, HW, AW and MLP. Asklepios Biobank as part of the DZL is partly funded by BMBF.

## Extended data figures

**Supplementary Figure 1.**
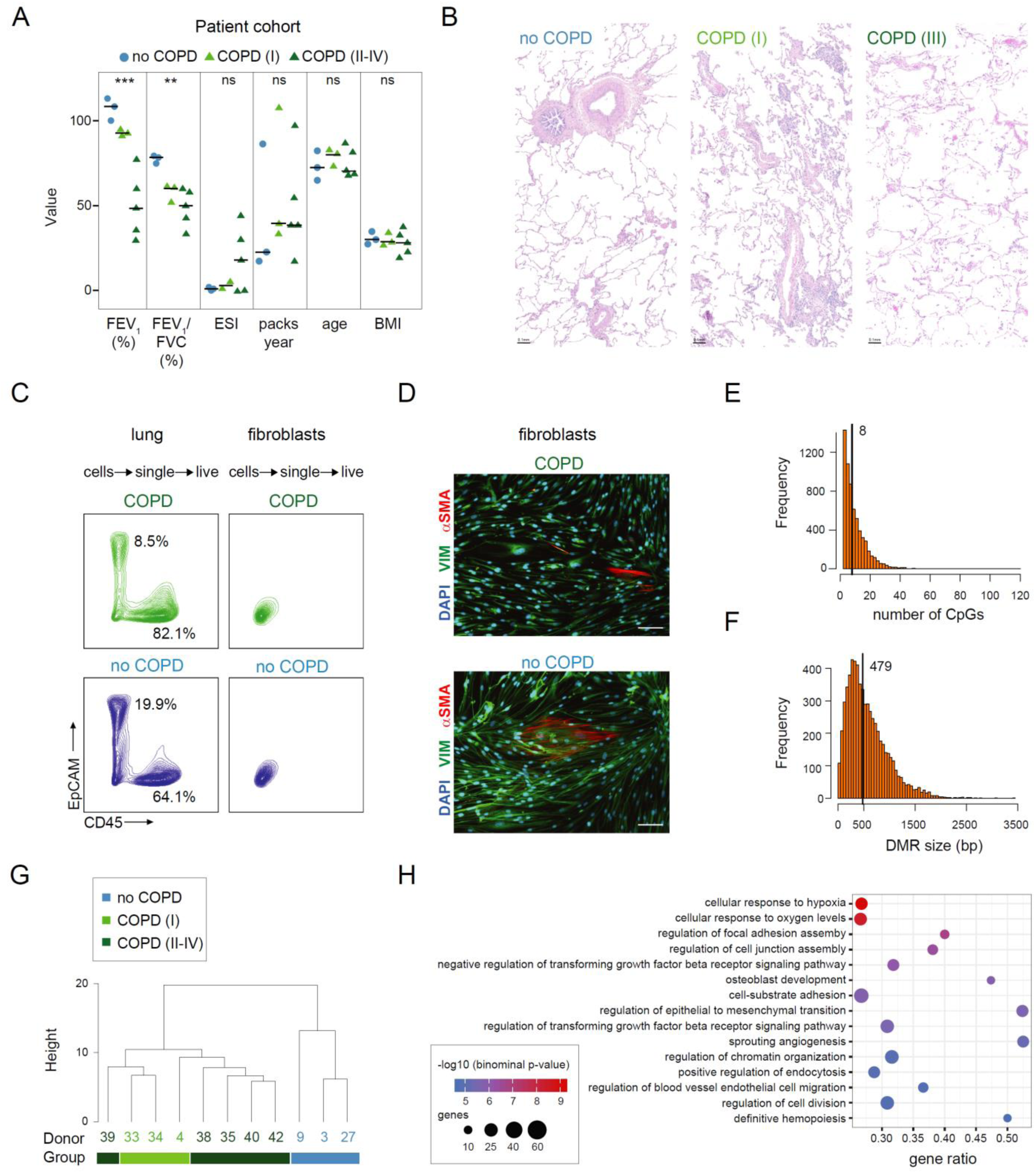
Genome-wide DNA methylation changes occur early in human lung fibroblasts in COPD and progress with disease development (supporting information Figure 1) **A** Characteristics of the lung tissue donors used in this study for fibroblast isolation (BMI, body mass index; FEV_1_, forced expiratory volume in 1 s; FVC, forced vital capacity; ESI, emphysema score index). Lung function between COPD and smoker control group (no COPD) is significantly different (*p-value < 0.05; non-parametric unpaired Kruskal-Wallis test). **B** Examples of hematoxylin and eosin (H&E) stainings of no COPD, COPD (I) and COPD (II-IV) tissue samples. **C-D** Validation of the purity of isolated fibroblast by FACS (**C**) and immunofluorescence (**D**). **C** Left, contour plot of lung live-single cell suspensions from COPD (green) and no COPD (blue) donors gated as indicated above. Right, contour plot of trypsinized fibroblasts from COPD (green) and no COPD (blue) donors. EpCAM-PE was used as an epithelial marker and CD45-Bv605 as an immune marker. **D** Immunofluorescence staining of fibroblasts isolated from the lungs of COPD (top) and no COPD (bottom) donors and stained with antibodies against vimentin (VIM, mesenchymal marker) and alpha smooth-muscle actin (αSMA, myofibroblast marker). DAPI was used to counterstain the nuclei. Scale bar: 0.1mm. **E-F** Histograms showing the number of CpGs per DMR (**E**) and the DMR size distribution (**F**). Median values are indicated and highlighted by a vertical line. **G** Hierarchical clustering of all samples based on 6,279 DMRs identified between no COPD and COPD (II-IV). **H** Functional annotation of genes located next to hypomethylated DMRs using GREAT. Hits were sorted according to the binominal p-value and the top 15 hits are shown. The adjusted p-value is indicated by the color code and the number of DMR associated genes is indicated by the node size.

**Supplementary Figure 2.**
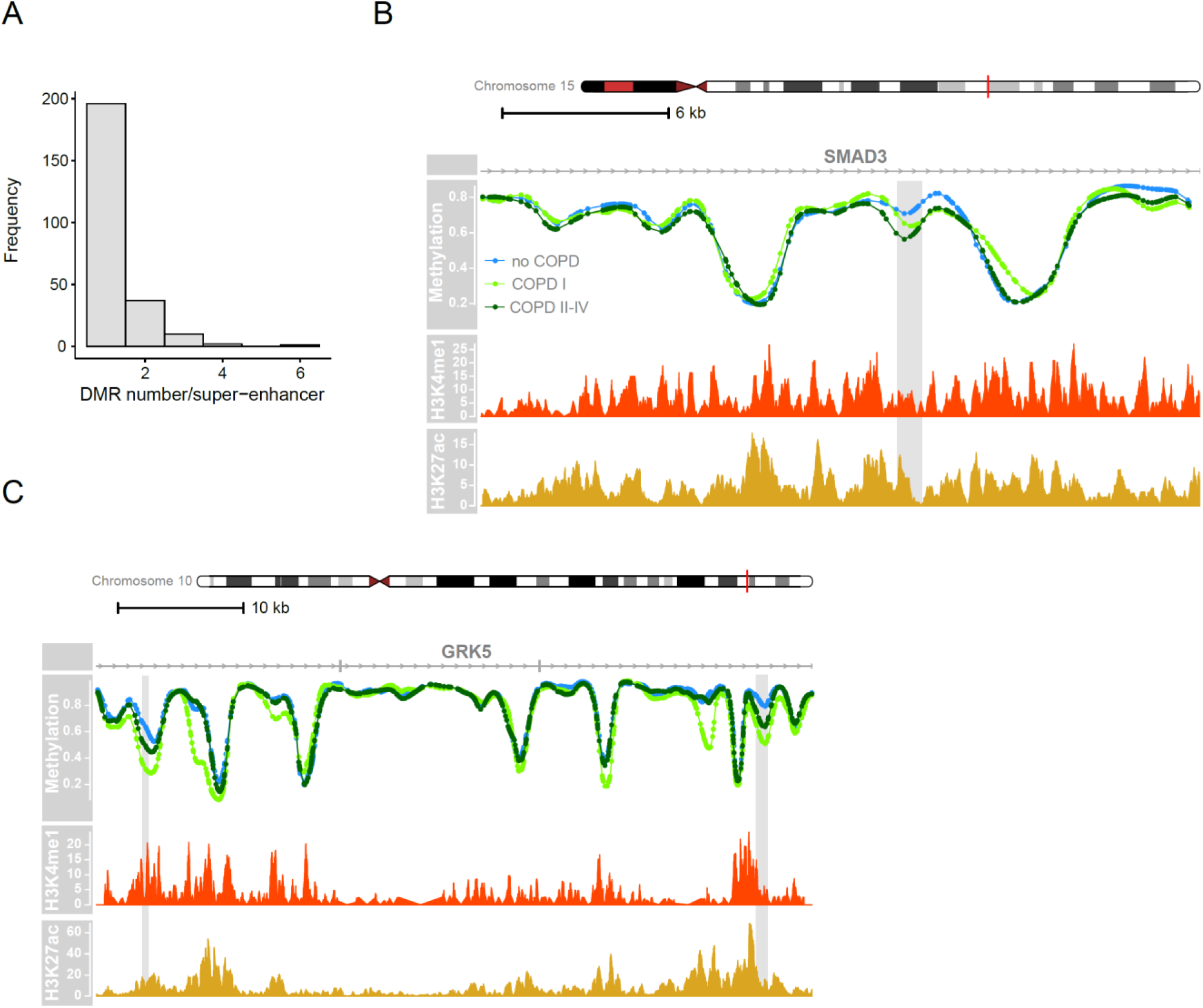
DNA methylation changes occur at regulatory regions in primary human lung fibroblasts cells during COPD (supporting information Figure 2) **A** Number of DMRs associated with super-enhancers **B-C** Genome browser views of two super-enhancer regions overlapping with identified DMRs (shaded in grey). Group median CpG methylation is shown for no COPD (blue), COPD (I) (light green) and COPD (II-IV) (dark green). At the bottom, the level of enhancer marks: H3K4me1 (ENCODE accession: ENCFF102BGI; red track) and H3K27ac (ENCODE accession: ENCFF386FDQ; orange track) is depicted as fold change over control.

**Supplementary Figure 3.**
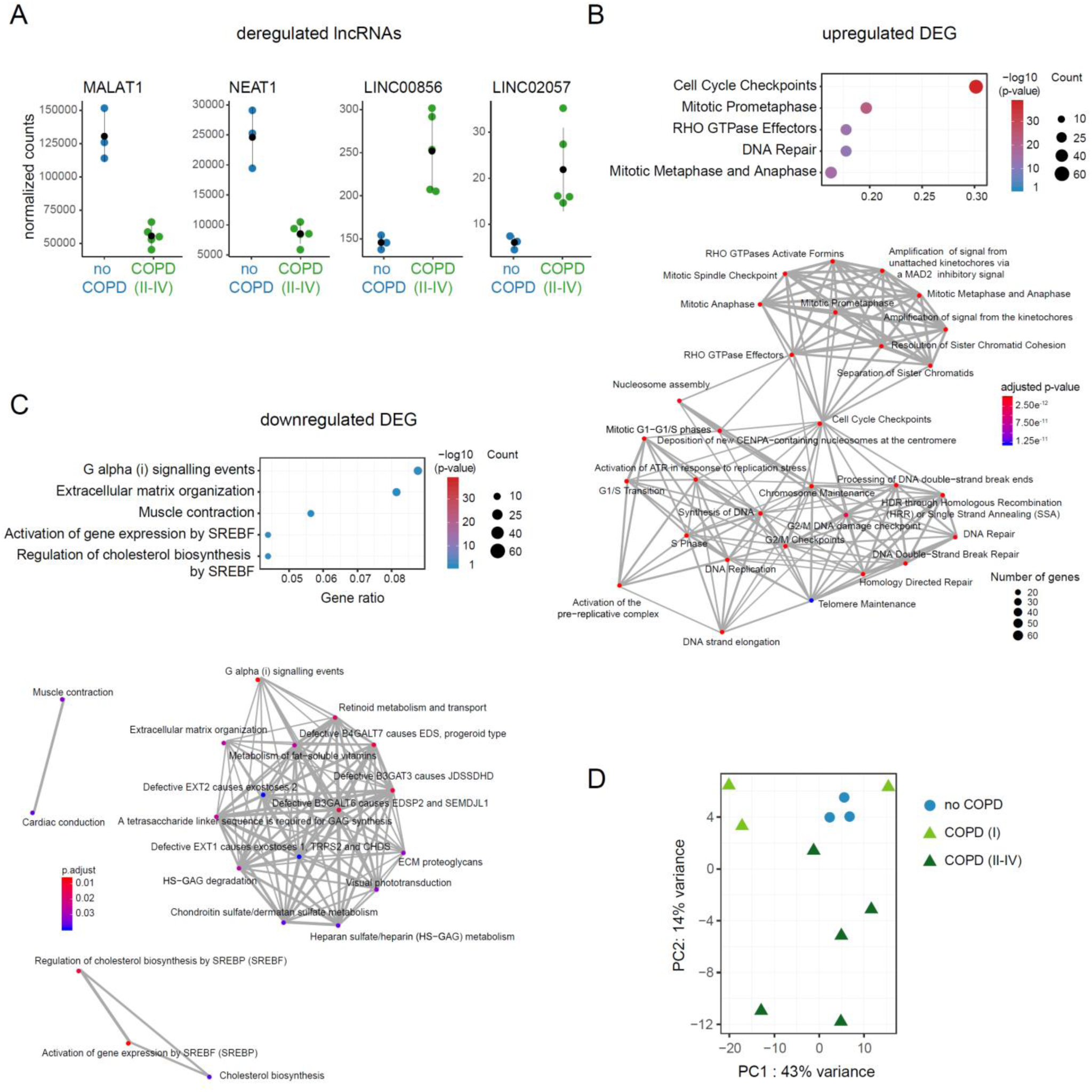
DNA methylation changes in primary human lung fibroblasts are accompanied by gene expression changes in COPD (supporting information Figure 3) **A** Normalized RNA-seq read counts of four exemplary differentially expressed lincRNAs **B-C** Functional annotation of upregulated (**B**) and downregulated (**C**) DEGs. Top 5 enriched GO biological process terms are shown on top of each panel. Bottom panels show a more detailed overview of the biological processes enriched for upregulated (**B**) and downregulated (**C**) DEGs. Biological processes are connected, if DEGs associated with both processes are in common. **(D)** Unsupervised principal component analysis (PCA) of all samples based on the 500 most variable expressed genes.

**Supplement Figure 4.**
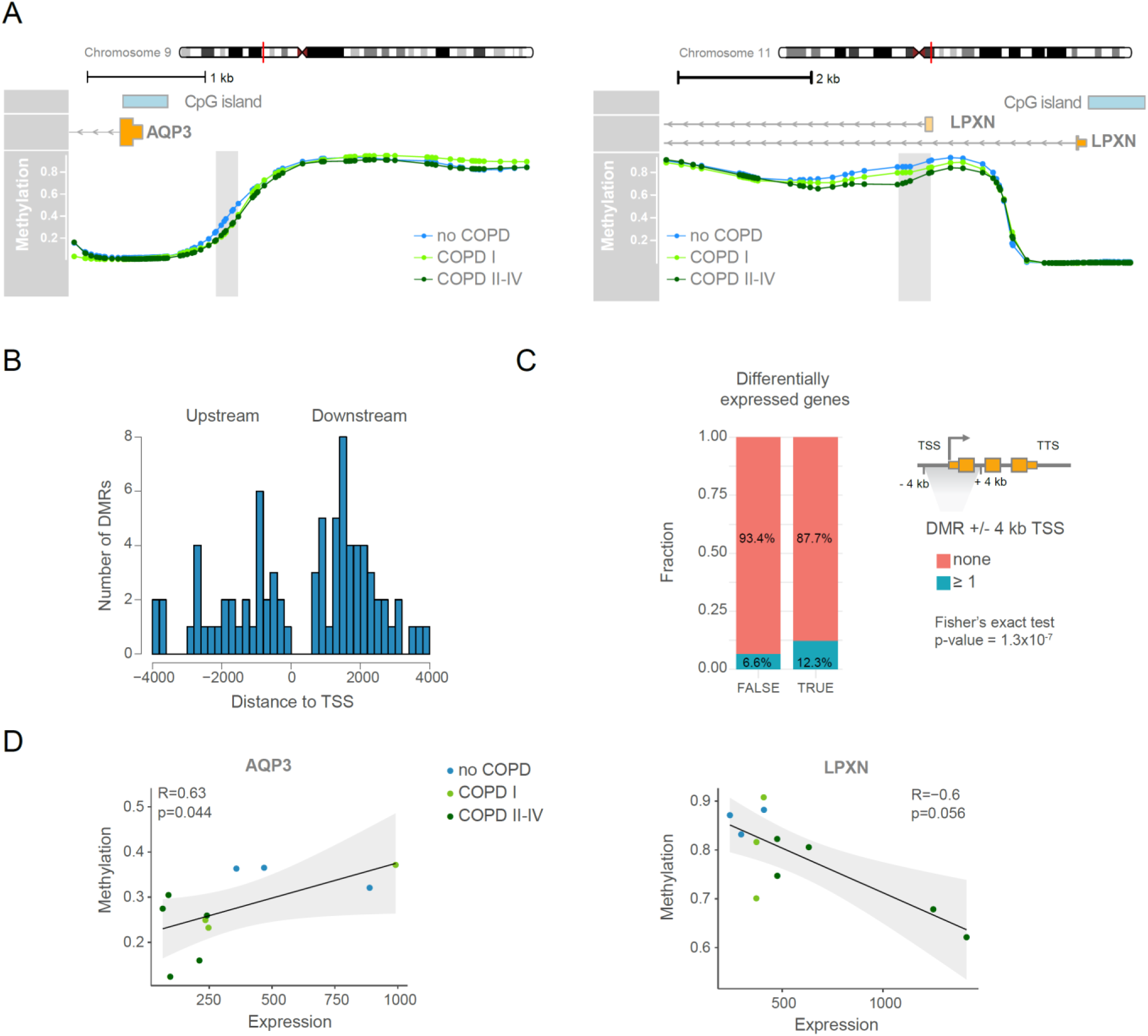
Integrative data analysis reveals epigenetically regulated genes in COPD fibroblasts (supporting information Figure 4). **A** Representative methylation profiles and DMRs (shaded in grey). Group median CpG methylation is shown for no COPD (blue), COPD (I) (light green) and COPD (II-IV) (dark green). RefSeq annotated genes and CpG islands are indicated. **B** Location of DMRs relative to the TSS of DEGs. **C** Fraction of genes associated with at least one DMR in the proximity (+/- 4kb) of the TSS are indicated in blue. Genes were split into not significantly changed genes (left bar, FALSE) and DEGs (right bar, TRUE). DMRs are significantly enriched at DEGs (Fisher’s exact test: p-value = 1.3×10^-7^). **D** Examples of genes showing correlation between gene expression and methylation. Scatter plots showing positive (left, AQP3) and negative (right, LPXN) correlation between gene expression and methylation of promoter-associated DMR. Each dot represents an individual donor. Dots are color coded according to disease state. Gene expression is illustrated as normalized counts. Methylation is illustrated as average beta value of the corresponding DMR. Linear regression analysis was performed (black line) and the 95% confidence interval is indicated (grey area). P-value and Spearman correlation coefficient (R) are indicated.

**Supplement Figure 5.**
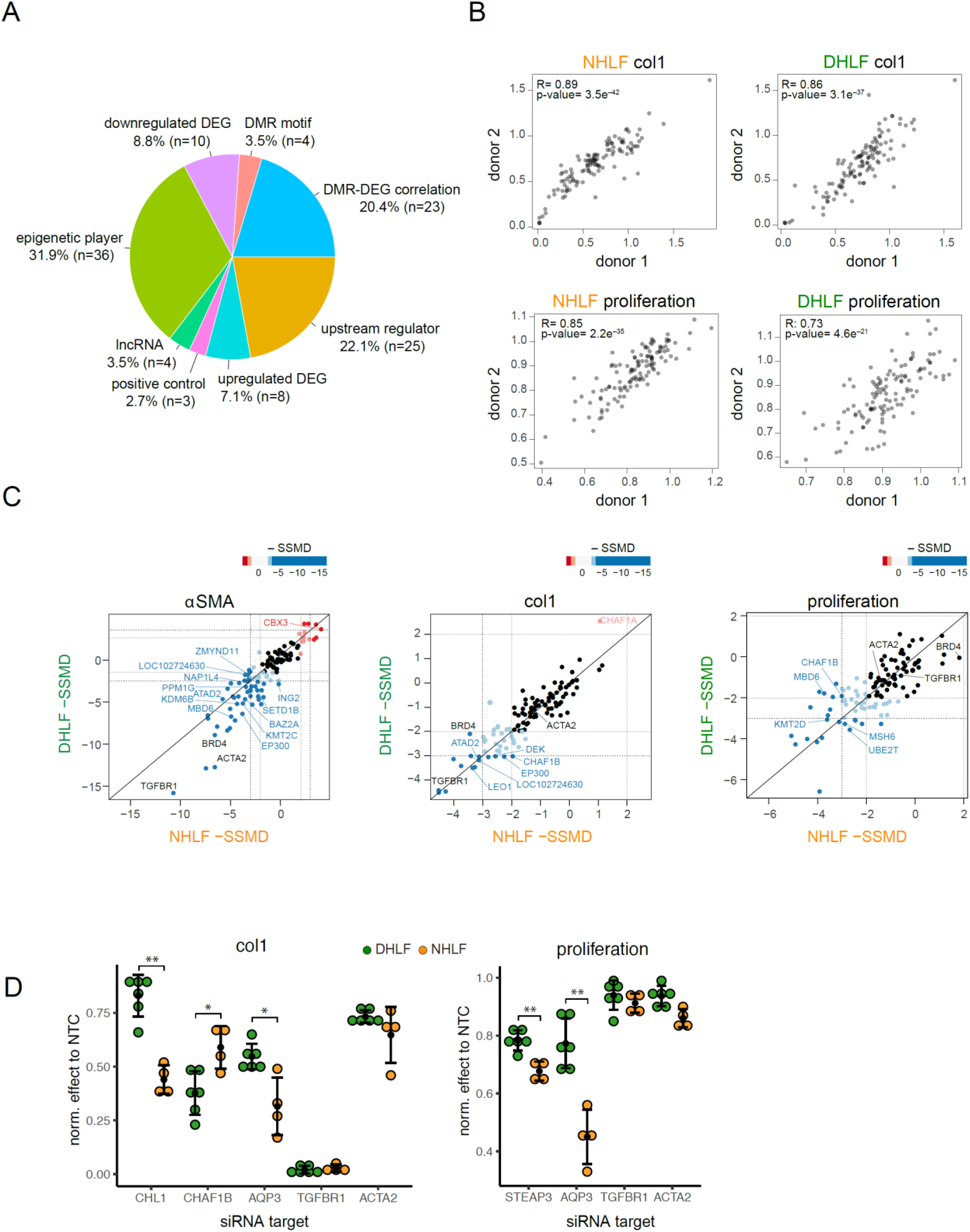
siRNA-based phenotypic screens in normal and COPD primary human lung fibroblasts identify multiple candidate genes regulating COPD phenotypes (supporting information Figure 5). **A** Distribution of candidates selected for functional validation among different categories (related to Figure 5C). The selected candidates were divided into 7 categories: (1) epigenetic players dysregulated on the transcriptional level, (2) DMR motif includes transcription factors with binding sites overrepresented in DMRs, (3) upstream regulators identified by the Ingenuity Pathway Analysis, (4) DMR-DEG correlation represents differentially expressed genes correlated with epigenetic changes in their promoter, (5) top downregulated genes, (6) top upregulated DEG, and (7) lncRNA for the top dysregulated long non-coding RNAs. In addition, 3 assay controls (ACTA2, TGFβR1 and BRD4) were included. **B** Scatterplots showing the correlation of the data obtained from 2 different NHLF (left) and 2 DHLF (right) donors in the phenotypic screens using col1 and proliferation/nuclei count as readouts. **C** Examples of epigenetic hits from the siRNA screen for each indicated readout. Each dot represents a unique candidate tested, blue and red dots represent significant hits (|SSMD values| ≥ 2 are shown in lighter shade and |SSMD values| ≥ 3 in stronger shade). Assay controls are labeled in black and epigenetic factors identified as strong positive hits (|SSMD values| ≥ 3) in fibroblast to myofibroblast transition and cell proliferation processes are labeled in blue or red depending on the effect on each readout upon KD. **D** Dotplots showing examples of positive hits with significant differences between NHLFs (2 donors, 2 biological replicates each, shown in green) and DHLFs (3 donors, 2 biological replicates each, shown in orange) in col1 and proliferation/nuclei count readout. Screen readout was normalized to the corresponding non targeting siRNA control (NTC). TGFβR1 and ACTA2 represent screen controls. The results of all replicates are shown. Statistical evaluation was performed with unpaired two-tailed student’s t-test. *: p-value < 0.05; **: p-value < 0.01. Black dots denote the means and error bars represent standard deviation.

